# The Duality of Insect Macroevolution: Pulsed Genomes and Gradual Morphology Shape Lineage Diversification

**DOI:** 10.1101/2025.11.05.686547

**Authors:** Akash Ajay, M. Sravya, K Akhila, K. Yashwant

## Abstract

Insects, comprising over half of all known species, are pivotal to ecosystems and ideal for studying trait evolution due to their morphological and genomic diversity. Yet, the macroevolutionary interplay between their phenotypic traits and genomic architecture is poorly resolved. Using a Tree of Life framework and phylogenetic comparative methods, we investigated the evolutionary dynamics of morphological and genomic traits across multiple insect orders with trait-specific taxonomic sampling optimized for data availability. Critically, while traditional metrics showed no correlation between species richness and evolutionary mode, the pattern of pulsed evolution revealed a clear macroevolutionary signal. Morphological diversification tended towards gradualism, whereas genomic evolution was predominantly pulsed. Notably, this pulsed signal manifested differently: large morphological pulses were associated with species-poor clades like *Odonata*, while species-rich clades exhibited major pulses in genome size. These results demonstrate that interpreting pulsed evolution’s contribution to diversification requires careful consideration of both trait identity and phylogenetic context. Further, for both morphological and genomic traits, the degree of phylogenetic conservatism was independent of the evolutionary distance between orders.

## 1. Introduction

Insects, being the taxonomic group with the largest number of species lineages (Misof et al., 2014), are a very diverse group of critical importance in ecology and economy (Losey and Vaughan, 2006; Cerritos, 2009). Insects have been one of the major biotic factors shaping the biosphere with co–co-evolutionary relationships with diverse taxa from plants to humans (Waage, 1979; Misof et al, 2014; Bruce, 2015; Joop and Vilcinskas, 2016). Insects stand out as one of the evolutionarily successful groups with high levels of speciation and low levels of extinctions (Engel, 2015; Schroer et al, 2018), adaptable body plans (Engel, 2015; Auman and Chipman, 2017), and evolution of social organization and behavior (Misof et al, 2014; Engel, 2015). Insects with their short generation times and easy culturing and storing have been sources of important model organisms (*Drosophila melanogaster*, *Apis mellifera*) and emerging model organisms (*Bombyx mori*, *Tribolium castaneum*) in biotechnology research (Adamski et al., 2019; Li et al., 2019). The study of the factors driving the niche diversification of insects, their morphological evolution, and the genes that drive this evolution remains an active field of research.

Previous comparative studies on insect morphology have successfully linked trait variation to ecological factors like land use (Bui et al., 2020) and extinction risk (Hagge et al., 2021), while others have focused on detailed morphological descriptions (Herbert et al., 2025) or tested for biomechanical links, such as the dissociation of mandible shape from diet (Edel et al., 2024). At a macroevolutionary scale, studies have shown that early burst models and punctuated equilibrium-based models are unlikely to explain morphological evolution (Harmon et al., 2010) and highlighted methodological advances for improving phylogenetic inferences (Duncan et al., 2025). Gao and Wu (2020) demonstrated that early burst models frequently characterize the evolution of genomic traits, including genome size, genomic GC content, rRNA GC, and N-ARSC (Average Nitrogen Atoms per Residual Side Chain), in their comparative analysis.

Deciphering the macroevolution of insects requires analyzing traits that capture key aspects of their phenotypic diversity and are linked to core functions, such as locomotion and feeding. This morphological perspective must be complemented by an analysis of genomic architecture, as features at this level represent fundamental elements that can constrain or catalyze evolutionary change, providing a necessary genomic perspective on diversification. For this study, we focused on key morphological traits—head dimensions (width, height, length), thorax width, forewing length, body length, and bite force—compiled from established datasets (Hagge et al., 2021; Staton et al., 2023; Ruhr et al., 2024). These traits were selected because they are fundamental to insect ecology and performance (Dudley, 2002; Ospina-Garces et al., 2018), directly influencing feeding mechanics (bite force, head dimensions), locomotion (thorax and wing dimensions), and overall body plan. By analyzing these traits across seven diverse insect orders (*Blattodea, Coleoptera, Diptera, Hymenoptera, Lepidoptera, Odonata,* and *Phasmatodea*) within a Tree of Life framework, this research aims to decipher the extent to which their evolution is shaped by phylogenetic history versus adaptive processes.

The selected genomic features—genome size, GC content, GC skew (Arakawa & Tomita, 2007), coding genes, non-coding genes, and chromosome number—serve as key indicators of broader evolutionary patterns. These whole-genome and karyotype characteristics offer a window into fundamental processes such as spontaneous mutational dynamics (Wang & Obbard, 2023; Lynch et al., 2023; Ajay et al., 2024), genome evolution and its structural aspects (Petro, 2001; Graphodatsky et al., 2011; Gao & Wu, 2022), as well as constraints and opportunities underlying morphological and organismal complexity (Mattick, 2001; Gregory, 2002; Choi et al., 2020; Ajay et al., 2024). This dual focus on morphological and genomic traits enables a comprehensive analysis of the evolutionary mechanisms driving insect diversification.

Following Edel et al. (2024)’s findings on phylogenetic signal in mechanical advantage potential in *Orthoptera*, we test whether bite force shows comparable phylogenetic signal and Brownian Motion across diverse insects. We further examine its integration with other morphological/genomic traits and assess pulses in its evolutionary trajectory. The research investigates the extent to which phylogeny shapes the evolution of the above-mentioned morphological and genomic traits, the relationships among them, and whether these traits are under the regime of phylogenetic conservatism or adaptive divergence —specifically testing if phylogenetic proximity among orders predicts shared evolutionary patterns. Additionally, the study examines the role of species diversity in shaping phylogenetic signals and the trait evolution models that govern genomic and morphological parameters. It examines whether these models indicate pulsed evolution (Hunt, 2008), characterized by discrete jumps, and quantifies the number, magnitude, and frequency of jumps per unit of branch length to identify patterns of rapid evolutionary change (Gao & Wu, 2022). By employing phylogenetic signal analyses, evolutionary model fitting, and phylogenetically corrected correlations, this study aims to elucidate the evolutionary processes driving insect diversity and address gaps in understanding the tempo and mode of trait evolution in this highly diverse lineage.

## 2. Methods

### 2.1 Data collection

#### 2.1.1 Morphological Features

Morphological data, encompassing head width, head height, head length, thorax width, forewing length, body length, and bite force, were obtained from Ruhr et al. (2024) for seven insect orders: *Blattodea, Coleoptera, Hymenoptera, Mantodea, Odonata, Orthoptera,* and *Phasmatodea*. Hagge et al. (2021) and Staton et al. (2023) also provided data for *Coleoptera,* including three of these features: head height, head width, and wing length. Due to methodological differences in trait measurement protocols across datasets (e.g., variations in landmark definitions, measurement tools, and sex-specific sampling), morphological trait data were analyzed separately for each source. This conservative approach minimizes potential bias in phylogenetic signal estimation and model fitting while preserving data integrity. Data for males and females were maintained and analyzed in separate datasets to mitigate the confounding effects of sexual dimorphism on trait evolution. For each species across all datasets, we calculated the mean morphological trait values, separating them by sex when data permitted. Orders with insufficient sampling (n < 10) were excluded from morphological analyses, including *Neuroptera*, *Megaloptera*, *Raphidioptera*, *Dermaptera*, *Embioptera*, and *Mantophasmatodea*. Four families with sufficient sampling (n > 10) - *Acrididae (Orthoptera)*, *Formicidae (Hymenoptera)*, *Mantidae (Mantodea)*, and *Carabidae (Coleoptera)* - were selected for cross-scale evolutionary comparisons to enable analysis of evolutionary patterns at different taxonomic scales. Supplementary Table 1 (ST1) presents morphological data at both order and family levels.

#### 2.1.2 Genomic Features

Genomic feature data, including genome size, GC content, coding genes, non-coding genes, and chromosome number for the followig orders - *Orthoptera, Blattodea, Odonata, Phasmatodea, Hemiptera, Coleoptera, Lepidoptera, Diptera, Hymenoptera*; were retrieved from the National Center for Biotechnology Information (NCBI) database. Data extraction from NCBI was facilitated using custom Linux scripts to automate the collection and compilation of genomic information across the target orders. The missing taxonomic information, if not available from the source databases, was filled with the help of the *taxsize* R package. GC skew was computed for each genome with the help of custom R scripts using the formulae (G-C) / (G+C). The genome features and accompanying taxonomy of the species are arranged order-wise with taxonomic information in Supplementary Table 2 (ST2).

#### 2.1.3 Species Diversity

Data on the total number of described species for each insect order and family analyzed in this study were gathered from established literature sources. This information on clade-specific diversity is presented in Supplementary Table 3 (ST3).

### 2.2 Building the Phylogenetic Framework

Phylogenies for each order were constructed using the Tree of Life framework, leveraging publicly available phylogenetic data to establish evolutionary relationships and enable phylogenetically informed analyses of trait evolution. Timetrees with their divergence time estimates are frequently used for analyzing the tempo and modes of trait evolution, particularly when assessing pulsed versus gradual evolutionary patterns (Wang &Obbard, 2023; Edel et al., 2024).

For the morphological analysis, we constructed a backbone phylogeny from a Newick tree (generated using the Tree of Life server) containing 13 insect orders -*Blattodea, Coleoptera, Dermaptera, Embioptera, Hymenoptera, Mantodea, Mantophasmatodea, Megaloptera, Neuroptera, Odonata, Orthoptera, Phasmatodea,* and *Raphidioptera*. Using a custom R script and the ape package, we identified tips via string and genus-level pattern matching, pruned the tree, and verified order assignments. A final backbone tree was generated by extracting the most recent common ancestor (MRCA) for each order, selecting one representative tip, and renaming the tips to match the order names for visualization with ggtree. For the genomic features analysis, an analogous approach was applied to a separate Newick tree spanning 22 orders: *Archaeognatha, Blattodea, Coleoptera, Dermaptera, Dictyoptera, Diptera, Ephemeroptera, Lepidoptera, Mantodea, Mecoptera, Megaloptera, Neolepidoptera, Neuroptera, Odonata, Orthoptera, Phasmatodea, Plecoptera, Siphonaptera, Strepsiptera, Thysanoptera, Trichoptera, and Zygentoma*. Enhanced matching incorporated expanded genera lists and fuzzy searches for common genomic model species to maximize coverage, with subsequent MRCA extraction and tip pruning yielding a comparable backbone structure. Both trees were rooted at the MRCA of all included orders, visualized with branch lengths in millions of years, and exported as Newick files for downstream comparative analyses.

The phylogenetic scope of this study is illustrated in Figure 1, detailing the relationships among the insect orders analyzed for (a) morphological traits and (b) genomic features. Figures 2 and 3 show the timetrees constructed for insect orders studied for morphological evolution and genomic evolution, respectively. Due to their large size, the comprehensive trees for *Diptera* and *Hymenoptera* are provided in the Zenodo repository [10.5281/zenodo.17524136]. Time-calibrated phylogenetic trees for the morphological evolution analysis are provided in Supplementary Figure 1 (SF1). For the genomic evolution analysis, trees for all families are available in the Zenodo repository [10.5281/zenodo.17524136], with representative families shown in Supplementary Figure 2 (SF2).

**Figure 1.**
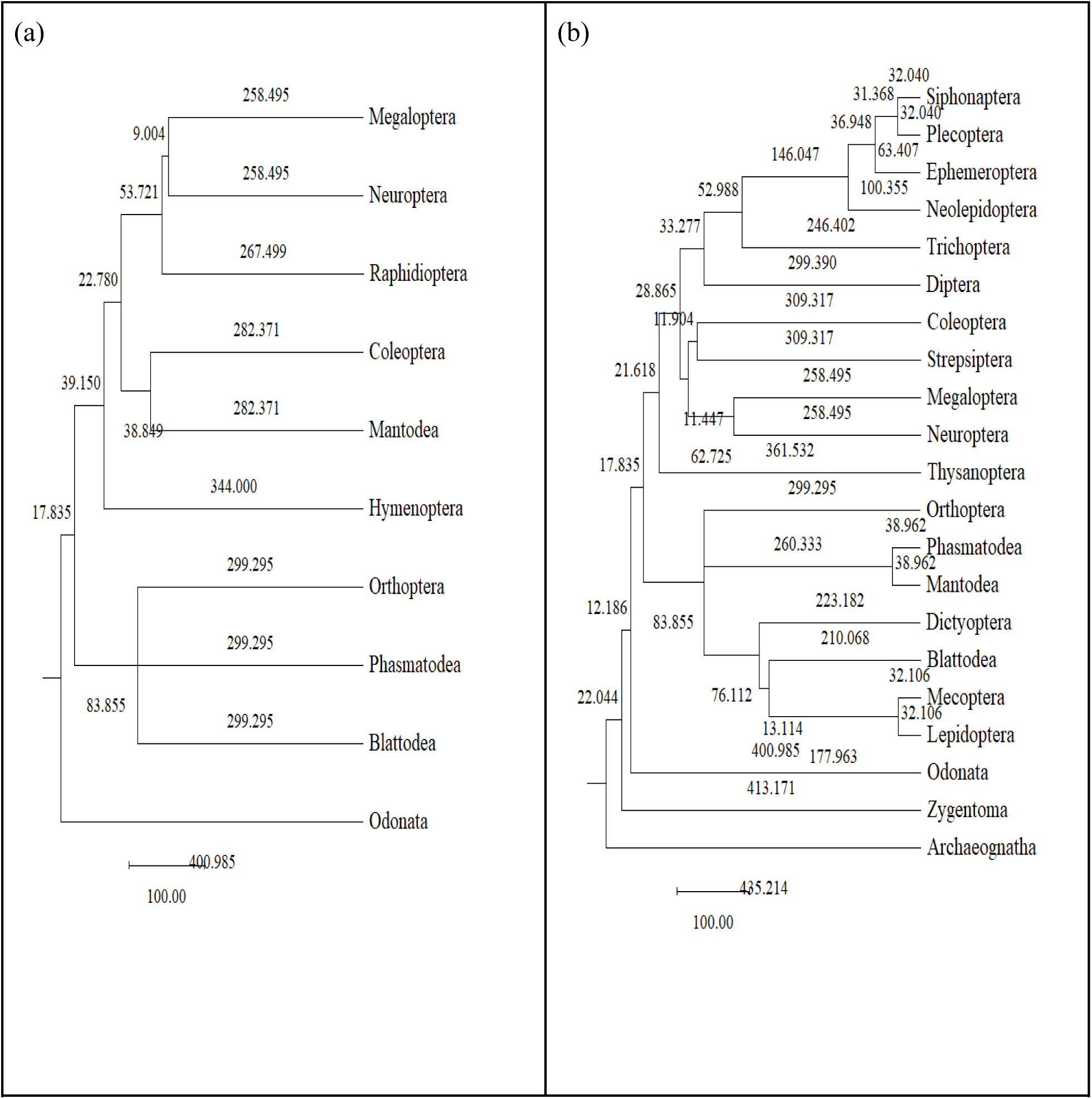
Phylogenetic frameworks for comparative analyses. (a) The backbone phylogeny used for analyzing the evolution of morphological traits. (b) The backbone phylogeny used for analyzing the evolution of genomic traits. Tip labels indicate the insect orders included in each analysis.

**Figure 2.**
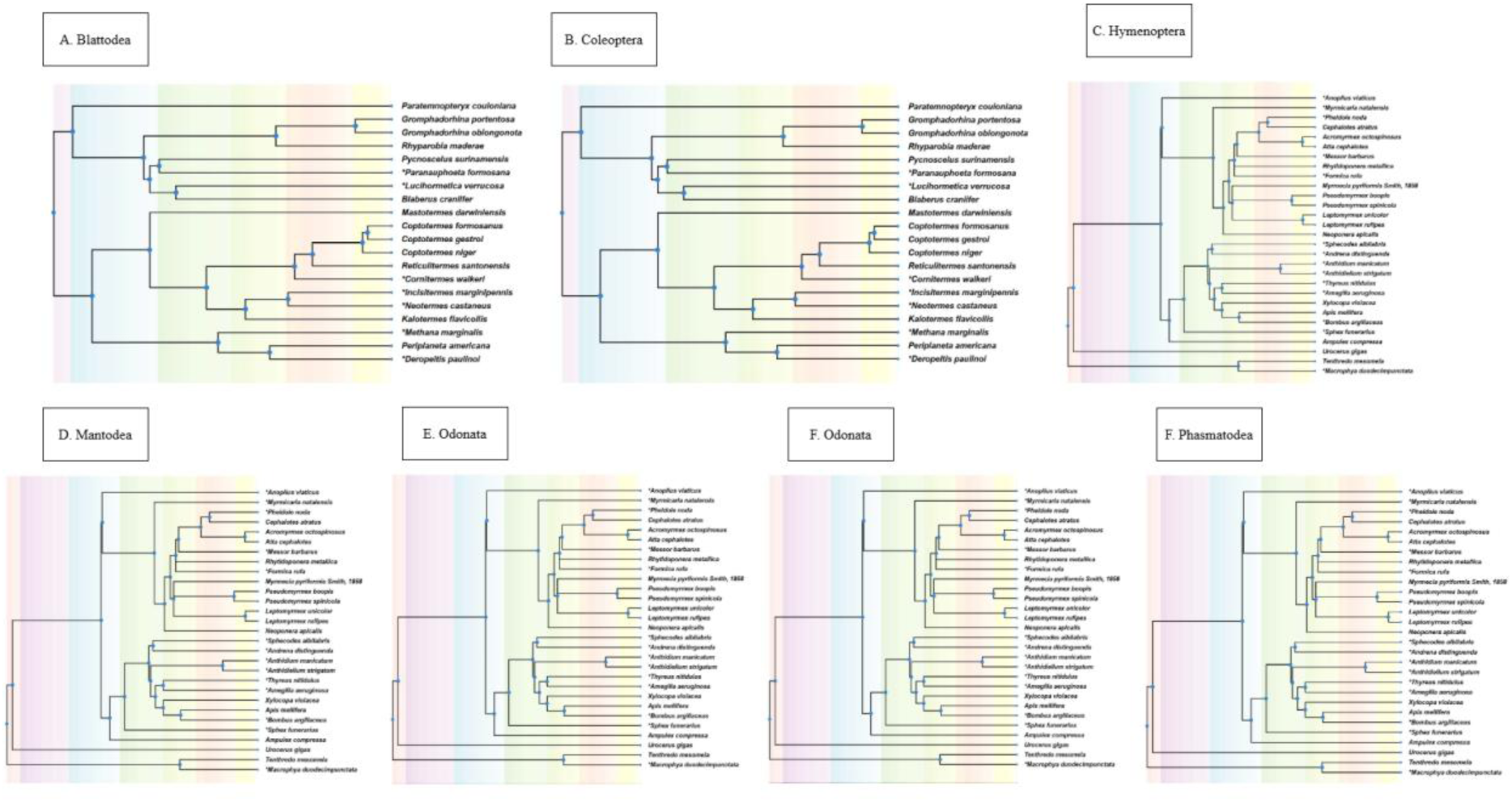
The timetree used for comparative analyses of morphological trait evolution. The tree includes the following orders: Blattodea, Coleoptera, Hymenoptera, Mantodea, Odonata, Orthoptera, and Phasmatodea.

**Figure 3.**
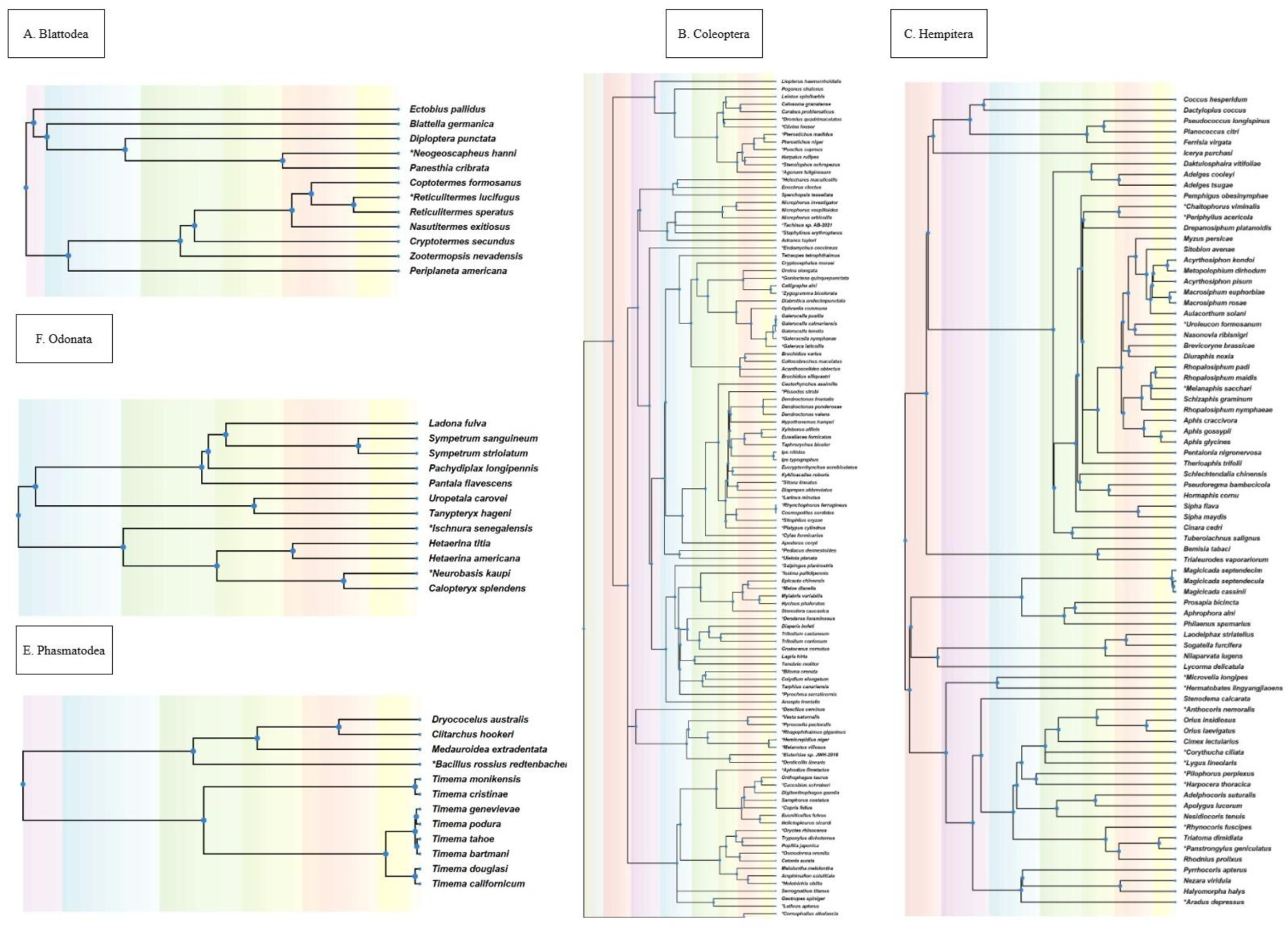
The timetree used for comparative analyses of genomic trait evolution. The tree includes the following orders: Blattodea, Coleoptera, Diptera, Hemiptera, Hymenoptera, Lepidoptera, Odonata, and Phasmatodea.

### 2.3 Exploratory Statistical Analysis

We used R (v4.3.2) to conduct exploratory analyses investigating morphological trait variation across seven insect orders and genomic trait variation across ten insect orders. One-way analysis of variance (ANOVA) was used to test for differences in trait distributions across orders, followed by Fisher’s exact test to assess significant differences in trait distributions among orders, providing an exploratory overview of trait variation for morphological traits (Supplementary Table 4) and genomic traits (Supplementary Table 5). Supplementary Figures 1 and 2 show the distribution of (a) morphological and (b) genomic trait values across the studied insect orders, visualized using violin plots with corresponding ANOVA results.

### 2.4 Phylogenetic Comparative Analyses

Phylogenetic comparative analyses were conducted primarily at the *order* level to ensure adequate sample sizes (≥20 species per analysis) and capture macroevolutionary patterns across Insecta. To validate order-level patterns and explore finer-scale dynamics, *family*-level analyses were conducted as supplementary analyses for families with adequate representation (≥10 species each). We first resolved phylogenetic polytomies using minimal branch length corrections (1×10⁻⁶) to ensure computational stability (Paradis et al., 2019). Phylogenetic signal was then quantified using Pagel’s λ (Pagel, 1999; Münkemüller et al., 2012) and Blomberg’s K (Blomberg et al., 2003; Münkemüller et al., 2012), implemented with custom R scripts leveraging the ape (Paradis et al., 2019) and phytools (Revell, 2024) packages to assess trait conservatism (Molina-Venegas et al., 2017). The trait models were fit with Brownian motion (BM) (Felsenstein, 1985), Orstein-Uhlenbeck (OU) (Hansen, 1997), Early-Burst (EB) (Harmon et al., 2010), Kappa (κ) (Pagel, 1999), Delta (δ) (Pagel, 1999; Pagel et al., 2006), and white noise (Harmon et al., 2015) models using the *geiger* package (Pennell et al., 2014). Akaike Information Criterion (Akaike et al., 2025) was used to estimate the best-fit model.

The phylogenetic signal (λ and K) for each trait was tested for correlation with the species richness of its respective order to identify any significant relationships. Orders were categorized as species-rich or species-poor using a 50,000-species threshold. Associations between these categories and evolutionary models were tested with Fisher’s exact tests for both morphological and genomic traits.

To test whether evolutionary proximity predicts similar phylogenetic constraints, we analyzed the relationship between phylogenetic distance and phylogenetic signal (λ) similarity across insect orders. For both morphological and genomic datasets, we calculated pairwise phylogenetic distances from order-level backbone phylogenies. We then performed linear regressions for each trait, testing if the phylogenetic distance between orders predicted the absolute difference in their λ values.

We analyzed phylogenetically corrected trait correlations using two complementary approaches. Initially, a broad-scale screening of pairwise relationships was conducted using Phylogenetic Generalized Least Squares (PGLS) (Symond & Blomberg, 2014), implemented in the *caper* package (Orne et al., 2013). To then obtain robust, multivariate inferences on the key morphological factors influencing our focal traits, we implemented phylogenetic mixed models in the *phyr* package for bite force and wing length. This framework was selected for its ability to generate more reliable parameter estimates by explicitly modeling phylogenetic non-independence as a random effect and accounting for intraspecific variation

### 2.5 Testing and fitting Pulsated Evolution for Insect Features

To test for pulses in trait evolution, we applied complementary statistical approaches to assess whether phylogenetically independent contrasts (PICs) (Garland et al., 1992) significantly deviate from the distributions expected under standard evolutionary models. Following the framework of Wu & Gao (2022), who tested PIC distributions against Brownian Motion (BM) expectations, we expanded this approach to include comparisons against Ornstein-Uhlenbeck (OU) models. We employed both the Kolmogorov-Smirnov (KS) test (Berger et al., 2014) and Anderson-Darling (AD) test (Scholz and Stephen, 1986) (using the *nortest* package (Gross et al., 2015)), with the latter providing enhanced sensitivity to deviations in distribution tails—where signals of evolutionary pulses manifest as outlier contrasts. The complementary nature of KS and AD tests strengthens inference against neutral and constrained trait distributions—while KS detects general distributional deviations, AD offers heightened sensitivity to the tail anomalies characteristic of evolutionary pulses.

Significant deviations from normality, as indicated by the KS test and AD test, supported the exploration of punctuated models (PE1, PE1+BM, PE2, PE2+BM, PE3+BM) for traits showing non-Brownian or non-OU patterns (Gao & Wu, 2022). Trait evolution was modeled using a methodological approach adapted from Gao & Wu (2022) - Brownian Motion (BM); one-, two-, and three-rate Poisson jump processes (PE1, PE2, PE3); and hybrid models combining each jump process with a background BM component (PE1+BM, PE2+BM, PE3+BM). In this parameterization, σ² represents the rate of BM evolution, while λᵢ and σ²ᵢ define the rate and variance, respectively, of the *i*-th evolutionary jump class. Model performance was evaluated using the Akaike Information Criterion (AIC), and the parameters of the best-supported model for each trait were analyzed to elucidate evolutionary patterns. Model selection between Brownian Motion and pulsed evolution variants was assessed using ΔAIC criteria from Burnham & Anderson (2002). Following their recommendations, we considered |ΔAIC| < 2 as substantial support for both models, ΔAIC = 4-7 as considerably less support for the higher-AIC model, and |ΔAIC| > 10 as essentially no support. This conservative approach ensured robust inference of evolutionary modes. These models were fitted using custom R scripts based on the *diversitree* package (FitzJohn, 2012). Figure 4 illustrates the analytical framework for phylogenetic analyses conducted in this study.

**Figure 4:**
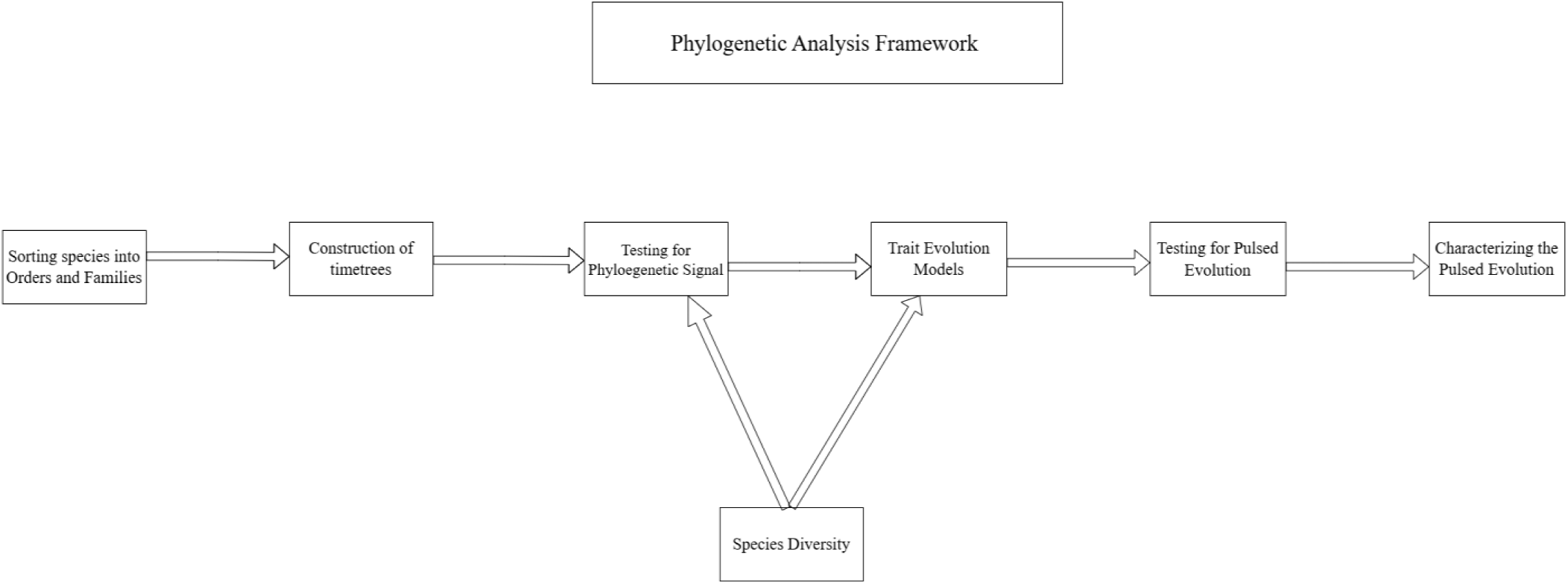
Analytical framework for phylogenetic analyses conducted in this study

## 3. Results

Analysis of variance revealed significant differences in all six morphological traits across the six insect orders (P < 2.23E-06), confirming substantial inter-order variation. Distinct morphological profiles emerged: *Mantodea* and *Odonata* consistently exhibited the largest head widths and forewing lengths, while *Hymenoptera* were among the smallest in these traits. *Orthoptera* and *Mantodea* had the greatest head height, whereas *Phasmatodea* showed notably larger head length and thorax width. *Blattodea* displayed the highest bite force, significantly exceeding all other insect orders studied.

Analysis of variance revealed significant differences in all genomic features across insect orders (P ≤ 0.0145), underscoring substantial genomic divergence. Distinct patterns emerged: *Hemiptera* and *Trichoptera* possessed the largest genomes, while *Hymenoptera* had the smallest. *Hymenoptera* and *Diptera* exhibited the highest GC content, in contrast to the lower values in *Hemiptera* and *Trichoptera*. *Hemiptera* and *Coleoptera* contained the highest number of coding genes, whereas *Hymenoptera* had the fewest. *Diptera* featured the highest number of non-coding genes. Chromosome numbers varied dramatically, with *Lepidoptera* having the highest count and Diptera the lowest. Further details are available in SF 1 & 2 and ST 4 & 5.

### 3.1 Testing for Phylogenetic Signal -

#### A. Morphological Traits

Phylogenetic signal analysis across seven insect orders—*Blattodea, Coleoptera, Hymenoptera, Mantodea, Odonata, Orthoptera,* and *Phasmatodea*—revealed varying degrees of evolutionary conservatism in morphological traits: head width, head height, head length, thorax width, wing length, bite force, and body length (*Mantodea, Orthoptera,* and *Odonata* only) (Supplementary Table 6 (ST6)). *Coleoptera, Orthoptera,* and *Phasmatodea* exhibited strong, significant phylogenetic signal across nearly all traits (e.g., *Orthoptera* head width, λ = 0.99, *P* < 0.05; *Phasmatodea* head length, K = 1.32, *P* = 0.001). This finding for *Coleoptera* is further supported by two other independent datasets (Hagge et al., 2021; Staton et al., 2023), which also showed a strong signal for head and wing morphology (e.g., head length, λ = 0.93, P < 0.001). *Hymenoptera* and *Blattodea* showed a strong signal for most traits, though head length was a notable exception.

*Coleoptera* exhibited a strong signal for most traits in our primary analysis, a pattern nuanced by sex-specific data from Staton et al. (2023), which revealed stronger phylogenetic conservatism in female body size than in male traits. We observed a conflicting pattern for wing length in *Coleoptera* between datasets of Ruhr et al. (2024) and Staton et al. (2023). The signal was present in the broader phylogenetic context of Ruhr et al. (2024) but not in the smaller taxonomic group studied by Staton et al. (2023), indicating that patterns of trait conservatism can be scale-dependent. In *Mantodea* and *Odonata*, signal strength was more variable and often trait-specific, with λ and K values sometimes presenting conflicting results (e.g., *Odonata* head length, λ = 0.95, *P* < 0.001 but K = 0, *P* > 0.05). A consistent pattern emerged for bite force, which demonstrated no significant phylogenetic signal in any order (e.g., *Blattodea* bite force, λ ∼ 0, *P* = 1.0), indicating its evolution is highly decoupled from phylogeny.

At the family level, distinct patterns emerged: *Acrididae* (*Orthoptera*) exhibited consistently strong phylogenetic signal across all head dimensions and body size traits (head length: λ = 1.00, *P* = 0.001; K = 0.58, *P* = 0.001), while *Formicidae* (*Hymenoptera*) showed significant signal for bite force (λ = 1.05, *P* = 0.03; K = 1.20, *P* = 0.008) and thorax width but not for head dimensions (ST6). *Mantidae* demonstrated a generally weak phylogenetic signal, with most traits showing non-significant λ values, though head length displayed moderate conservatism (K = 1.00, *P* = 0.105). Figure 5 (a &b) presents heatmaps of phylogenetic signal across insect orders, showing Pagel’s λ (a) and Blomberg’s K (b) values for morphological traits, with significance indicators distinguishing patterns of trait evolution.

**Figure 5.**
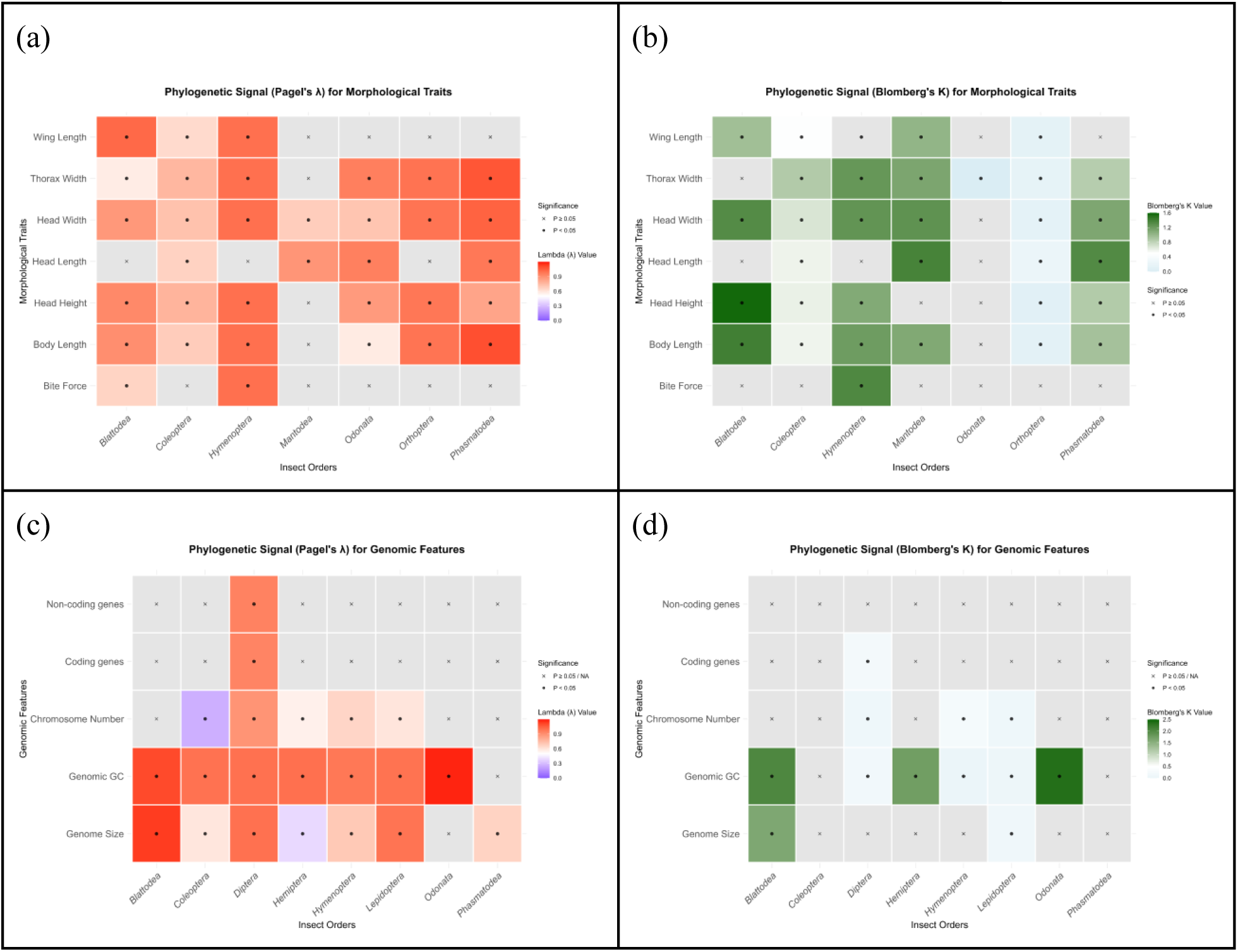
Phylogenetic signal heatmaps across insect orders for morphological (a-b) and genomic (c-d) traits. (a,c) Pagel’s λ values indicate phylogenetic signal strength relative to Brownian motion. (b,d) Blomberg’s K values measure trait conservatism. Solid circles denote significant signals (P < 0.05); crosses indicate non-significant results.

No significant correlation was found between species richness and morphological phylogenetic signal metrics (Pagel’s λ and Blomberg’s K) for any of the traits across the insect orders examined (Table 1). Our findings, though limited by the number of orders sampled, provide no evidence that species richness is a primary driver of phylogenetic trait conservatism. Expanding taxonomic sampling in future work will be crucial to test the generality of this pattern (Figure 5). A correlation analysis between species richness and phylogenetic signal was restricted to the order level, as the sample size of only four families was deemed insufficient to detect a reliable statistical relationship.

**Table 1.**
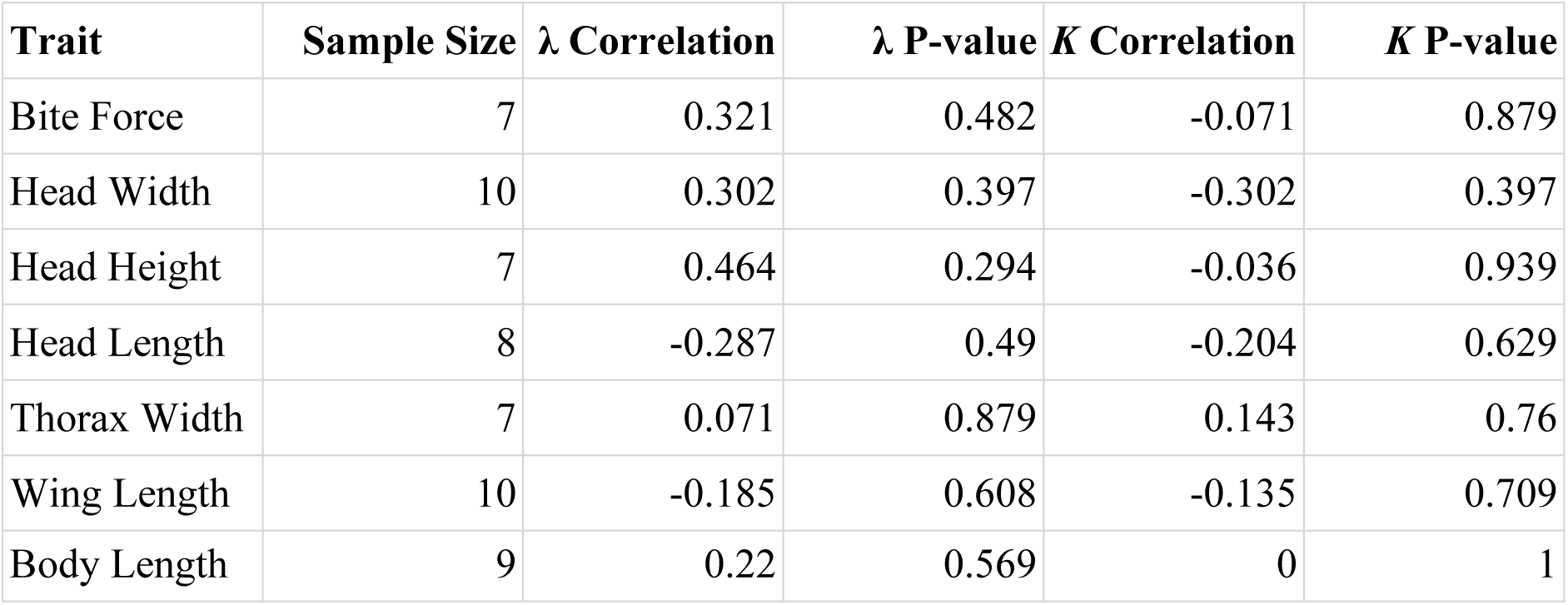
Correlations between phylogenetic signal (λ and Blomberg’s K) for morphological traits and lineage species richness

Analysis of individual morphological traits revealed no significant relationship between the phylogenetic distance of insect orders and the similarity of their evolutionary constraints. The relationship was non-significant across all measured traits (e.g., Body Length: R² = 0.1336, *P* = 0.1031, N_pairs = 21; Head Width: R² = 0.0564, *P* = 0.299, N_pairs = 21). This indicates that the degree of phylogenetic conservatism in morphological evolution is largely independent of the evolutionary relatedness between orders, though sample size limitations warrant caution (Supplementary Table 7 (ST7)).

#### Genomic Traits

Phylogenetic signal analysis of genomic traits revealed a clear hierarchy of evolutionary conservatism across insect orders (Supplementary Table 8 (ST8)). Core genomic architecture—including genome size and GC content—showed the strongest phylogenetic signal, being highly conserved in orders like *Blattodea* (λ > 1.10), *Diptera*, and *Lepidoptera* (λ > 0.99). Chromosome number displayed more variable evolutionary modes, with a significant signal in *Hymenoptera* and *Diptera* but weaker patterns in others. Analysis of GC skew revealed a stark dichotomy in its evolutionary mode across insect orders. *Diptera* (λ = 0.99, *P* < 1 E-58) and *Lepidoptera* (λ = 0.93, *P* < 1e-66) exhibited a strong and significant phylogenetic signal, indicating that GC skew is a conserved trait evolving in close association with phylogeny in these lineages. *Coleoptera* showed a weak but significant signal for Pagel’s λ (λ = 0.22, *P* = 0.045), suggesting a modest degree of phylogenetic structure. In contrast, *Hymenoptera* (N = 160) and *Hemiptera* (N = 70) both demonstrated a negligible and non-significant phylogenetic signal (λ ≈ 0), providing robust evidence that GC skew evolves independently of phylogeny in these two large and diverse orders. The similarly low signals in *Blattodea* and *Odonata* are more difficult to interpret definitively due to their smaller sample sizes. Similarly, coding and non-coding gene counts were overwhelmingly evolutionarily labile, showing no significant phylogenetic signal in any order where data were sufficient. This spectrum of conservation suggests that while fundamental genome structure is often phylogenetically constrained, finer-scale sequence composition and gene content evolve with greater evolutionary freedom. Figure 5 (c &d) presents heatmaps of phylogenetic signal across insect orders, showing Pagel’s λ (c) and Blomberg’s K (d) values for genomic traits, with significance indicators distinguishing patterns of trait evolution.

This pattern of strong phylogenetic conservatism in core genomic traits was largely reinforced at the family level (Table 3). Families such as *Nymphalidae* and *Drosophilidae* exhibited strong phylogenetic signal for genome size and GC content (λ > 0.92), mirroring the order-level trend. For chromosome number, the pattern was more heterogeneous. While a significant signal was detected in some families (e.g., *Apidae*, *Hesperiidae*), its absence in others like *Aphididae* (*Hemiptera*) and *Noctuidae* (*Lepidoptera*) could reflect genuine variability in evolutionary mode among constituent lineages of an order. However, the notably strong phylogenetic signal for chromosome number across the entire *Lepidoptera* order suggests that the lack of signal in specific families like *Noctuidae* (n=22) is more parsimoniously explained by insufficient statistical power at this taxonomic resolution than by a fundamental absence of phylogenetic signal across the order. Family-level analysis of GC skew largely reinforced the order-level findings. The strong phylogenetic signal observed across Lepidoptera was consistently recovered within its major constituent families, including *Nymphalidae*, *Hesperiidae*, and *Drosophilidae*. Conversely, the lack of phylogenetic signal observed in *Hymenoptera* and *Hemiptera* at the order level was also evident in their respective families, *Apidae* and *Aphididae*.

No consistent significant correlation was found between species richness and genomic phylogenetic signal metrics (Pagel’s λ and Blomberg’s K) across insect orders and families (Table 2). At the family level, all correlations were non-significant and often negative. However, this analysis was also constrained by the number of families with sufficient data (N_Families = 11-17) (Table 2, Supplementary Figure SF 3a & b). Collectively, these results provide no robust evidence that clade species richness is a general predictor of phylogenetic signal strength in genomic traits. However, more comprehensive sampling is required to draw definitive conclusions about these evolutionary patterns for both morphological and genomic traits.

**Table 2.**
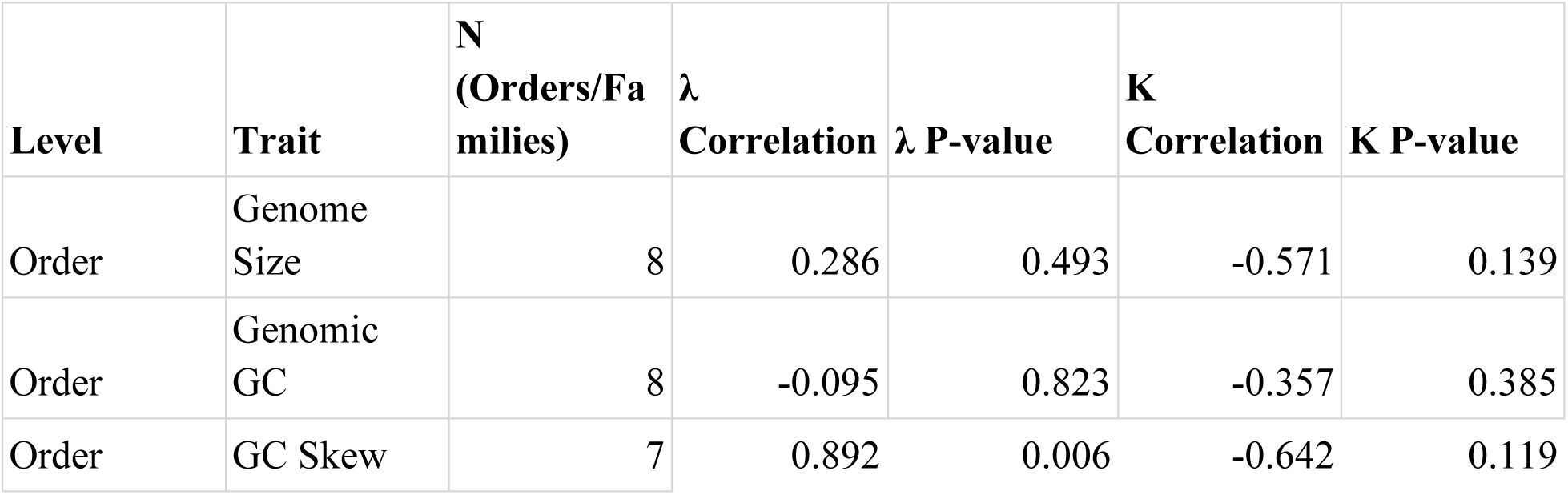
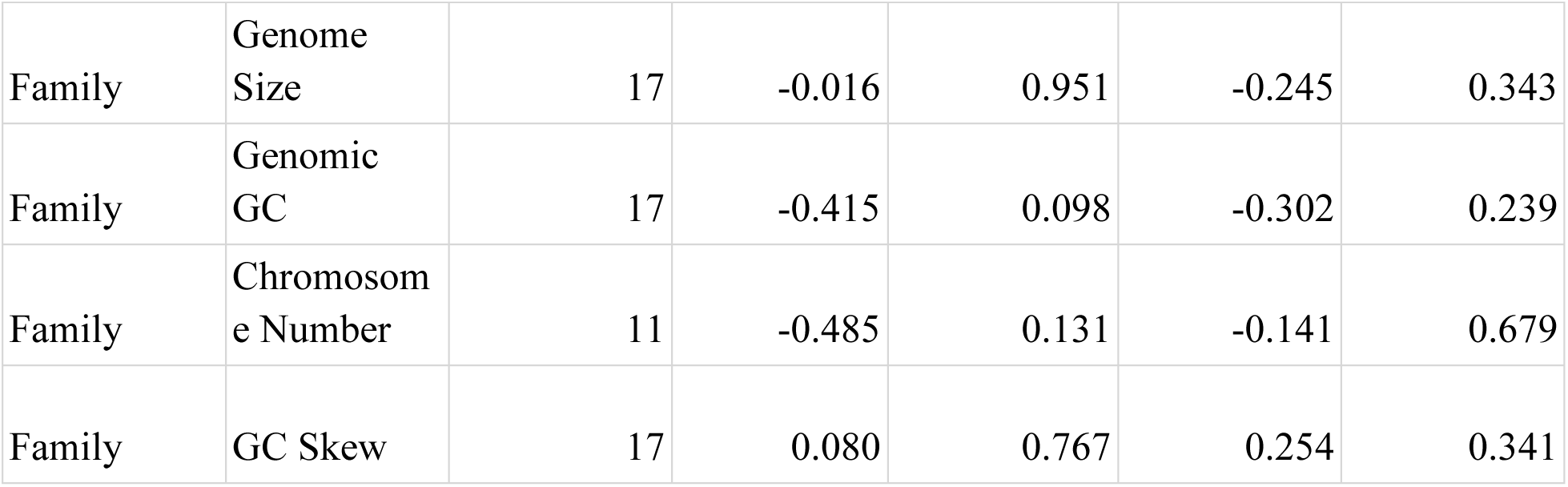
Correlations between phylogenetic signal (λ and Blomberg’s K) for genomic traits and lineage species richness

Analysis of genomic traits revealed that phylogenetic distance between orders does not predict similarity in evolutionary constraints, mirroring the pattern observed in morphological traits. Both genome size (R² = 0.004, *P* = 0.814, N_pairs = 15) and GC content (R² = 0.011, *P* = 0.713, N_pairs = 15) showed non-significant relationships with phylogenetic proximity (ST7). This indicates that the tempo and mode of genomic evolution are decoupled from the evolutionary relatedness of insect orders.

### 3.2 Trait Evolution Models

#### A. Morphological Traits

Order-specific patterns show that *Orthoptera* is characterized by pervasive stabilizing selection (OU) across nearly all traits, while *Coleoptera* exhibits a mix of OU processes and punctuated evolution (low kappa values) (Table 3). *Hymenoptera* displays a unique signature of neutral drift (Brownian Motion) for most traits, alongside an early-burst (EB) in wing evolution (a = −1.0E-06) . *Blattodea* and *Mantodea* are dominated by kappa models, indicating pulsed trait change. All traits in *Odonata* were best fit by a white noise model, indicating phylogenetic independent trait dispersion, a result likely attributable to low sample size [N_species =17] rather than a true biological signal (Table 3).

**Table 3.**
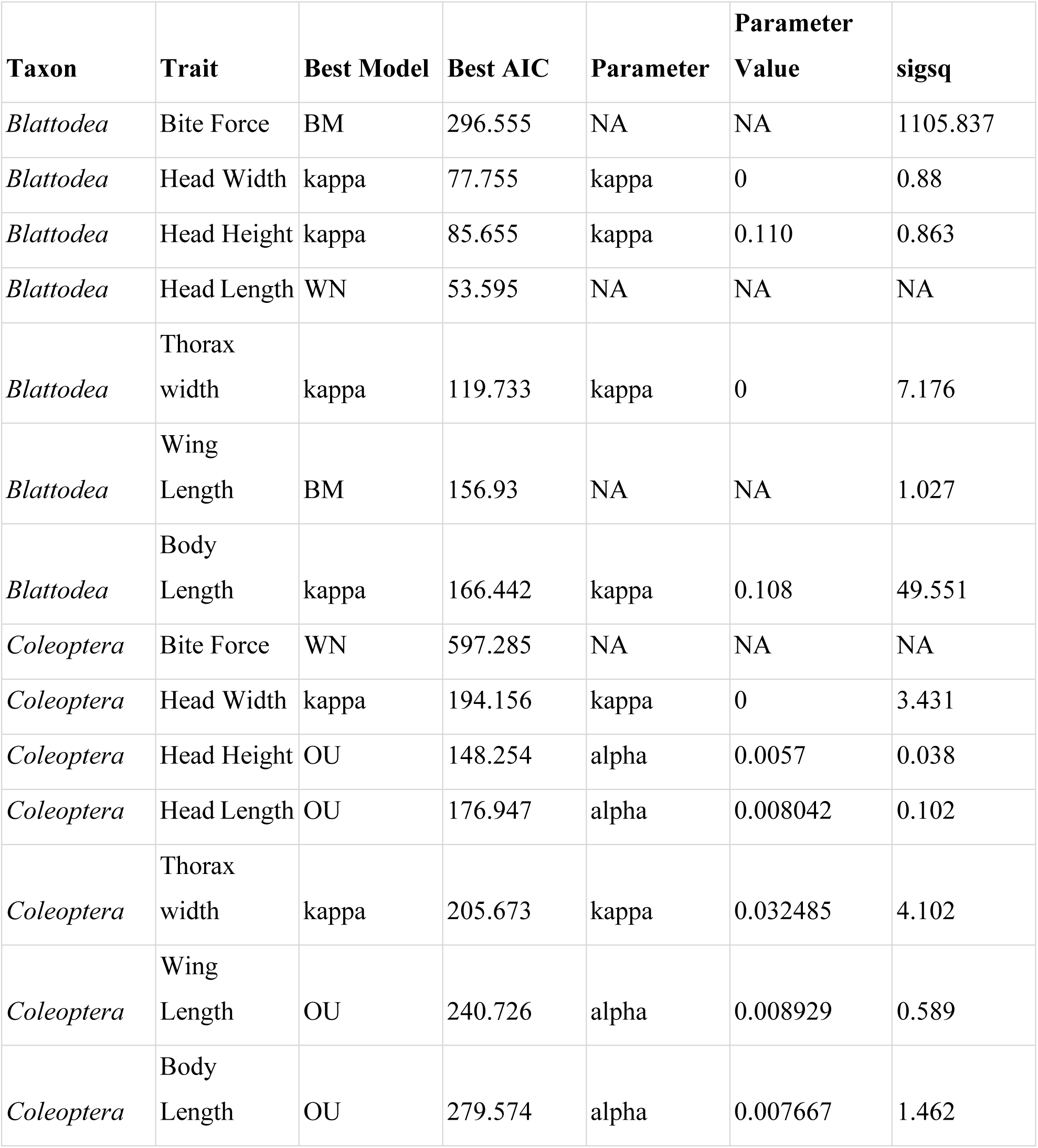
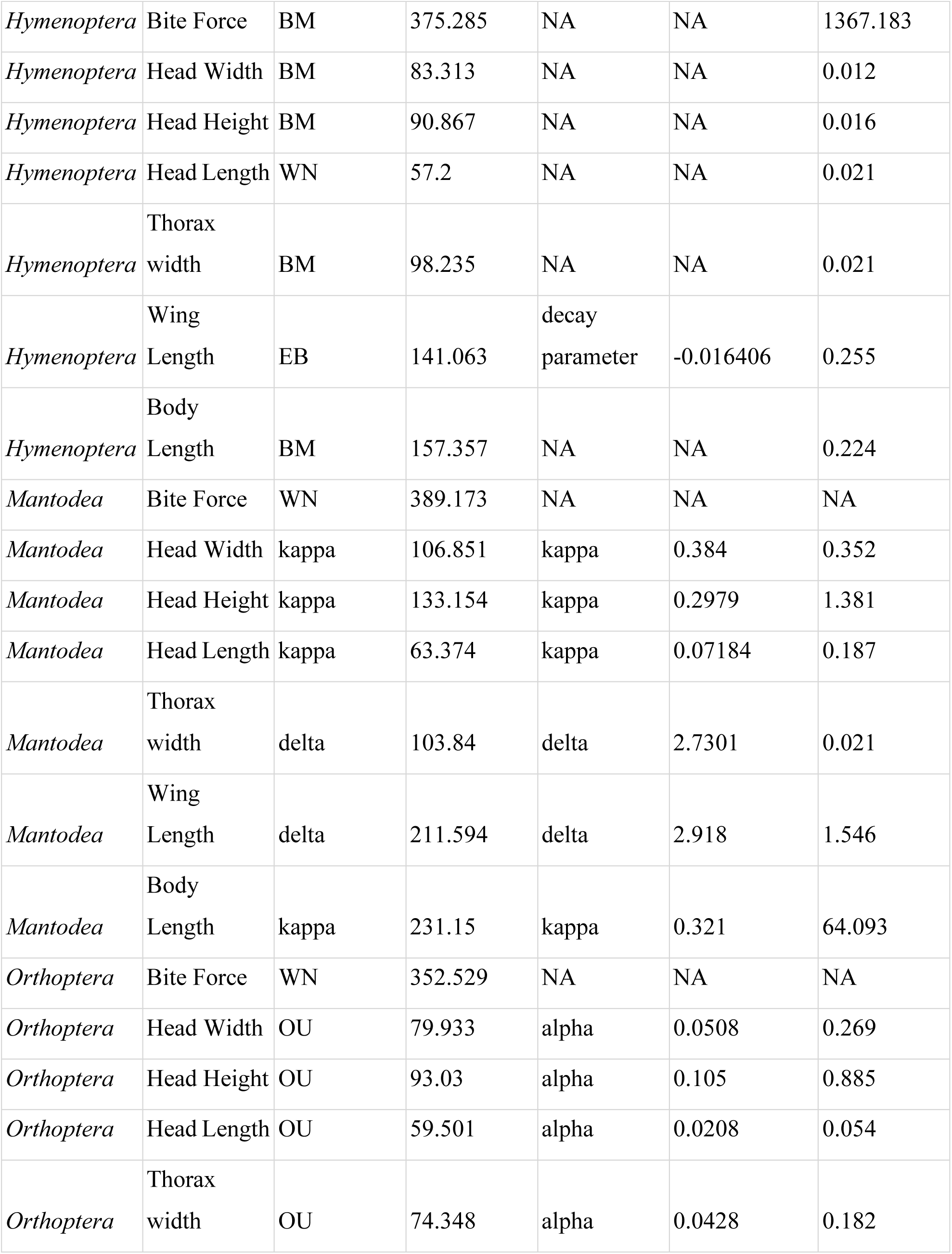
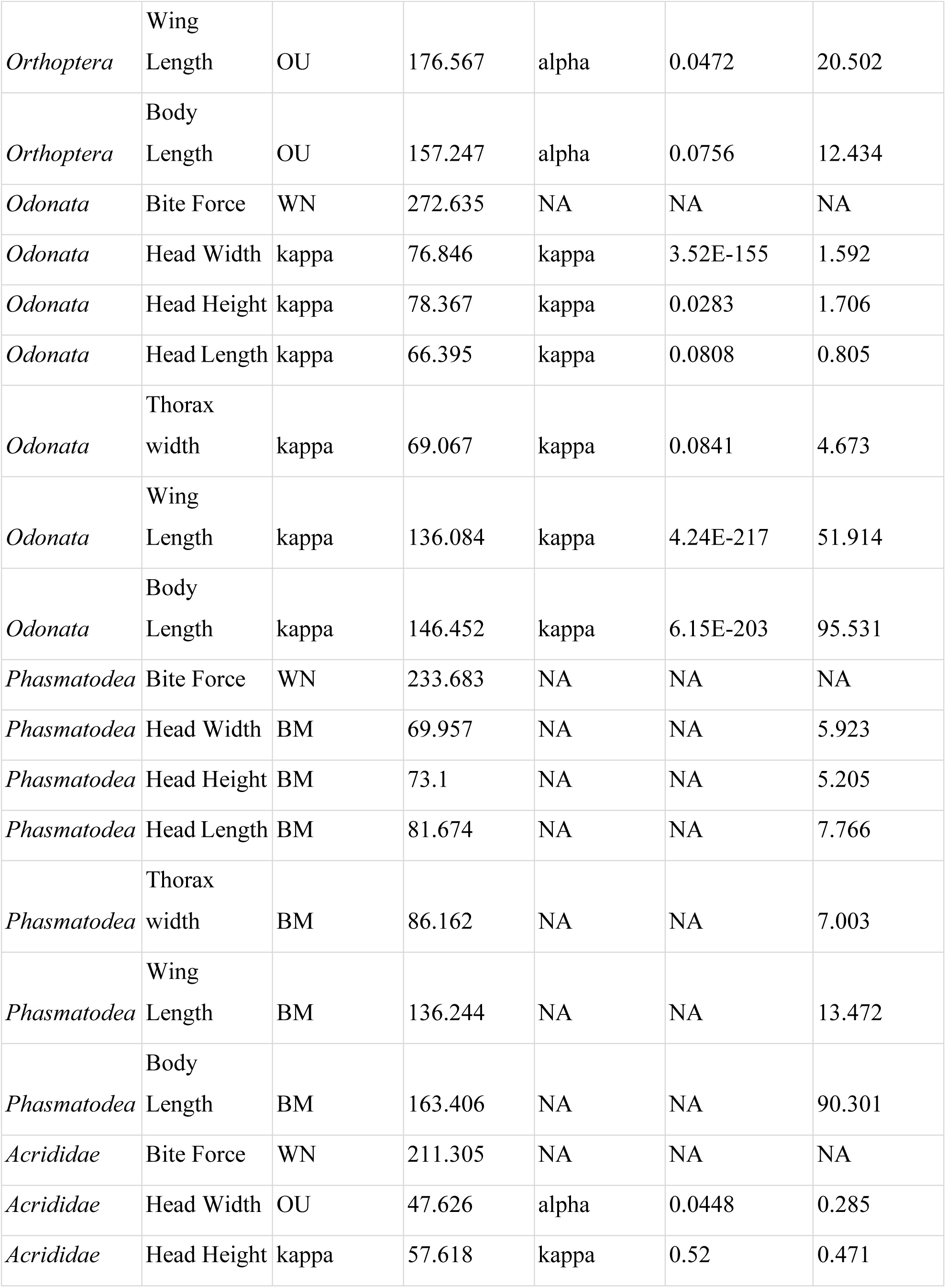
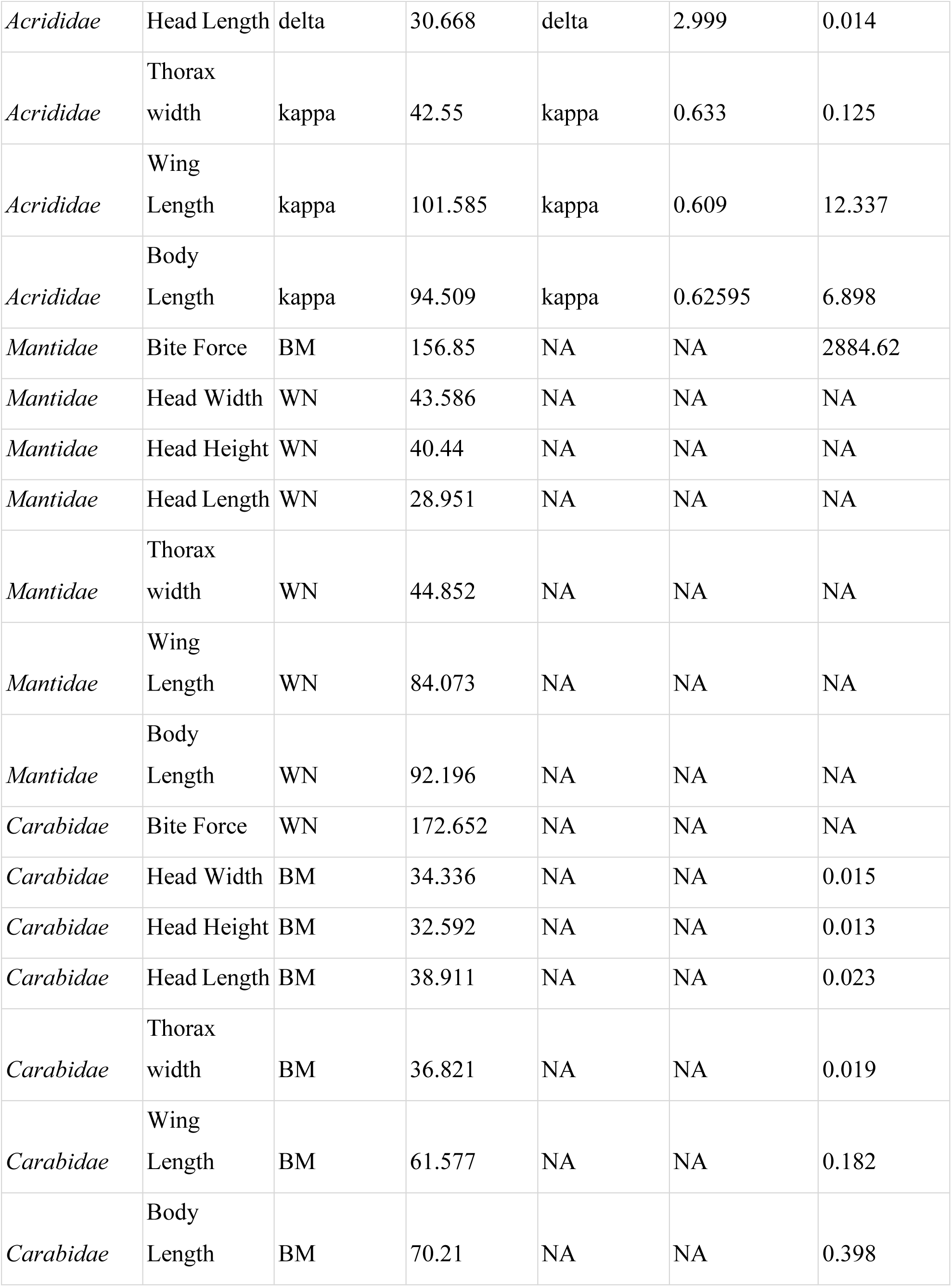
Model selection results and parameter estimates for the evolution of morphological traits across insect taxa. The best-fit model for each trait was selected using the Akaike Information Criterion (AIC) from a set of candidate models including Brownian motion (BM), Ornstein-Uhlenbeck (OU), early-burst (EB), and Pagel’s (kappa, delta) models. The relevant evolutionary rate (sigsq), constraint (alpha), or branch transformation (kappa, delta) parameter for the best model is provided.

Critically, trait-specific trends cut across these taxonomic patterns. Bite force consistently shows the highest evolutionary lability, being best fit by a white noise model in *Coleoptera*, *Mantodea*, and *Orthoptera*, indicating its evolution is largely decoupled from phylogeny in these lineages. In contrast, head dimensions most often evolve under constraint (OU or kappa). Independent datasets strongly support this pattern of constrained head evolution in *Coleoptera*; both the Hagge et al. (2021) and Staton et al. (2023) datasets found strong phylogenetic signal (Pagel’s Lambda) to be the best-fit model for head morphology, confirming this trait’s conserved evolutionary pattern in this order. The best-fitting model for each order and trait is detailed in Table 3, and the AIC values for all the models tested are provided in Supplementary Table 9 (ST9). These results demonstrate that an interplay between lineage-specific evolutionary modes and trait-specific functional constraints governs the macroevolution of complex morphologies. Figure 6 shows simulated morphological trait evolution in Coleoptera under best-fit models, with complete simulations for all orders available in Zenodo [10.5281/zenodo.17524136]

**Figure 6.**
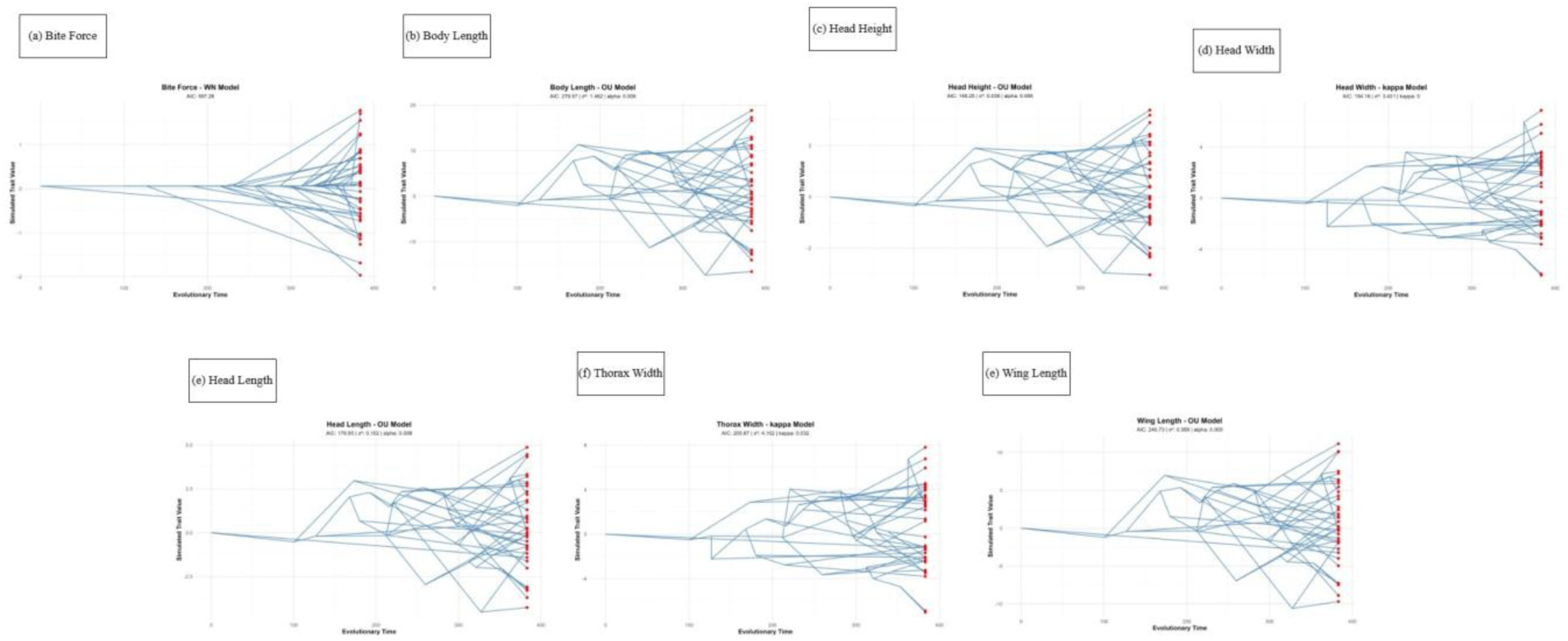
Simulated evolution of morphological traits in Coleoptera under best-fit evolutionary models

At the family level, the evolutionary patterns observed at the order level were largely maintained, though often with weaker support, likely due to reduced statistical power from smaller sample sizes. In *Orthoptera*, the family *Acrididae* reinforced the order’s trend of constrained evolution, with head and body traits best fit by Ornstein-Uhlenbeck (OU) and kappa models (e.g., head height, kappa AIC = 57.62). The high lability of bite force was also confirmed, with a white noise model being the best fit (AIC = 211.30). In *Hymenoptera*, the family *Formicida*e also showed a prevalence of neutral drift (Brownian Motion) for several traits, mirroring the order-level pattern. However, the best-fit models for head dimensions in *Formicidae* and for nearly all traits in *Mantidae* (*Mantodea*) and *Carabidae* (*Coleoptera*) were white noise, a result that strongly suggests insufficient phylogenetic signal detection due to limited within-family sampling rather than a true biological phenomenon (Table 3).

The evolutionary mode of morphological traits showed no significant relationship with species richness (Fisher’s exact tests: body length, *P* = 0.429; head height, *P* = 0.429; head width, *P* = 1.000; thorax width, *P* = 1.000; wing length, *P* = 0.810). This independence from diversification history, combined with their previously established phylogenetic conservatism, indicates that different macroevolutionary processes govern trait constraint versus lineage diversification.

Analysis of the best-fitting evolutionary models revealed no consistent association between a specific model of trait evolution (e.g., Brownian Motion, Ornstein-Uhlenbeck, Kappa) and the species richness of an order. The most species-rich orders—*Coleoptera*, *Hymenoptera*, and *Orthoptera*—each exhibited a different dominant evolutionary mode for their morphological traits. Evolutionary model comparisons for morphological traits, including AIC values for all fitted models, are provided in Supplementary Table 9 (ST9).

#### B. Genomic Traits

*Diptera* and *Lepidoptera* exhibited a consistent regime of punctuated evolution (Kappa) for core architecture, with genome size (*Lepidoptera*: κ=0.54, AIC=7,420) and chromosome number (Diptera: κ=0.11, AIC=290) evolving under the same pulsed dynamic (Table 4). In *Coleoptera*, genome size evolved under strong punctuated change (κ = 0.12, AIC = 1,675), yet chromosome number showed a weak phylogenetic signal, best fit by a White Noise model (AIC = 403.6). This discrepancy highlights their independent evolutionary histories and may be influenced by the lower sample size for chromosome data (N = 62 vs. N = 110 for genome size). In *Lepidoptera*, chromosome number evolution with kappa = 0 reduces to Brownian motion, representing neutral trait evolution. Furthermore, analyses of gene counts were likely confounded by undersampling, as seen in their frequent fit to White Noise models (e.g., *Coleoptera* non-coding genes, AIC=231.3; *Hemiptera* coding genes, AIC=487.6). The repeated finding of phylogeny-independent models in smaller clades (e.g., *Odonata, Phasmatodea*; N_species = 12) is likely an artifact of low statistical power, which diminishes the reliability of phylogenetic signal detection in these groups. Thus, while punctuated change is a major driver of genomic evolution in data-rich clades, evolutionary inferences for undersampled traits and orders should be interpreted with caution.

**Table 4.** Model selection results and parameter estimates for the evolution of genomic traits across insect taxa. The best-fit model for each trait was selected using the Akaike Information Criterion (AIC) from a set of candidate models including Brownian motion (BM), Ornstein-Uhlenbeck (OU), early-burst (EB), and Pagel’s (kappa, delta) models. The relevant evolutionary rate (sigsq), constraint (alpha), or branch transformation (kappa, delta) parameter for the best model is provided

| **Taxon** | **Trait** | **Best_Model** | **Best_AIC** | **Parameter** | **Parameter_**<br>**Value** | **sigsq** |
| --- | --- | --- | --- | --- | --- | --- |
| *Blattodea* | GC_skew | WN | -116.08176 | NA | NA | 3.57E-07 |
| *Blattodea* | Genome<br>Size | EB | 159.98499 | decay<br>parameter | -0.0291 | 56246.017 |
| *Blattodea* | Coding<br>genes | EB | 97.264 | decay<br>parameter | -0.054 | 1824621.63<br>2 |
| *Blattodea* | Genomic<br>GC | BM | 40.64492 | NA | NA | 0.0158 |
| *Blattodea* | Non -<br>coding<br>genes | BM | 98.966 | NA | NA | 54820.399 |
| *Coleoptera* | Genome<br>Size | kappa | 1675.125 | kappa | 0.118765 | 60918.839 |
| *Coleoptera* | Genomic<br>GC | kappa | 470.477 | kappa | 0.354741 | 0.470599 |
| *Coleoptera* | Non -<br>coding<br>genes | WN | 232.508 | NA | NA | 13851.867 |
| *Coleoptera* | Chromosom<br>e Number | WN | 405.574 | alpha | 0.794751 | 58.57339 |
| *Coleoptera* | Coding<br>genes | BM | 336.079 | NA | NA | 334620.115 |
| *Coleoptera* | GC_skew | OU | -753.35979 | alpha | 0.05470197 | 3.22E-07 |
| *Diptera* | Genome<br>Size | kappa | 4111.48 | kappa | 0.354165 | 2598.51 |
| *Diptera* | Genome GC | kappa | 1287.89 | kappa | 0.53183 | 0.404031 |
| *Diptera* | Non -<br>coding<br>genes | kappa | 1361.137 | kappa | 0.495959 | 503833.798 |
| *Diptera* | Chromosom<br>e Number | kappa | 290.533 | kappa | 0.114369 | 0.248949 |
| *Diptera* | Coding genes | kappa | 1372.966 | kappa | 0.595485 | 290976.278 |
| *Diptera* | GC_skew | kappa | -3124.8634 | kappa | 0.80293125 | 3.21E-07 |
| *Hemiptera* | Genome Size | kappa | 1216.317 | kappa | 0 | 234845.479 |
| *Hemiptera* | Genome GC | EB | 318.527 | decay parameter | -0.00924 | 314.233 |
| *Hemiptera* | Chromosome Number | kappa | 240.602 | kappa | 0.089697 | 4.501875 |
| *Hemiptera* | Coding genes | OU | 489.38 | alpha | 0.04298 | 2821984.836 |
| *Hemiptera* | GC_skew | WN | -769.41268 | NA | NA | 9.31E-07 |
| *Hemiptera* | Non - coding genes | WN | 362.2709 | NA | NA | 1502498.688 |
| *Hymenoptera* | Genome Size | kappa | 2348.784 | kappa | 0 | 22021.076 |
| *Hymenoptera* | Genome GC | kappa | 750.542 | kappa | 0.128279 | 1.369143 |
| *Hymenoptera* | Non - coding genes | OU | 802.93 | alpha | 0.270549 | 732286.085 |
| *Hymenoptera* | Chromosome Number | OU | 387.13 | alpha | 0.021713 | 1.444489 |
| *Hymenoptera* | Coding genes | OU | 883.766 | alpha | 0.026362 | 286096.741 |
| *Hymenoptera* | GC_skew | WN | -1487.5752 | NA | NA | 5.23E-06 |
| *Lepidoptera* | Genome Size | kappa | 7420.719 | kappa | 0.540379 | 2251.731 |
| *Lepidoptera* | Genome GC | kappa | 1708.471 | kappa | 0.424805 | 0.170868 |
| *Lepidoptera* | Non - coding genes | kappa | 671.186 | kappa | 0 | 942184.357 |
| *Lepidoptera* | Chromosome Number | kappa | 1998.044 | kappa | 0 | 13.465618 |
| *Lepidoptera* | Coding | OU | 834.84 | alpha | 1.82915 | 54644458.5 |
|  | genes |  |  |  |  |  |
| *Lepidoptera* | GC_skew | kappa | -6338.4425 | kappa | 0.48595895 | 1.34E-07 |
| *Odonata* | Genome Size | BM | 174.223 | NA | NA | 659.601 |
| *Odonata* | Genome GC | delta | 37.222 | delta | 0.258096 | 0.013582 |
| *Odonata* | Chromosome Number | BM | 23.993 | NA | NA | 0.010968 |
| *Odonata* | GC_skew | WN | -144.01167 | NA | NA | 2.58E-07 |
| *Phasmatodea* | Genome Size | kappa | 201.212 | kappa | 0 | 298253.008 |
| *Phasmatodea* | Genome GC | OU | 49.441 | alpha | 2.718282 | 11.885444 |
| *Phasmatodea* | Non - coding genes | BM | 72.965 | NA | NA | 22092.32 |
| *Phasmatodea* | Chromosome Number | BM | 42.626 | NA | NA | 0.129163 |
| *Phasmatodea* | Coding genes | BM | 84.338 | NA | NA | 379318.541 |
| *Phasmatodea* | GC_skew | WN | -81.664415 | NA | NA | 2.84E-07 |

Figure 7 displays genomic trait evolution in *Coleoptera* under best-fit models, while simulations for all other orders are archived in Zenodo [10.5281/zenodo.17524136].

**Figure 7.**
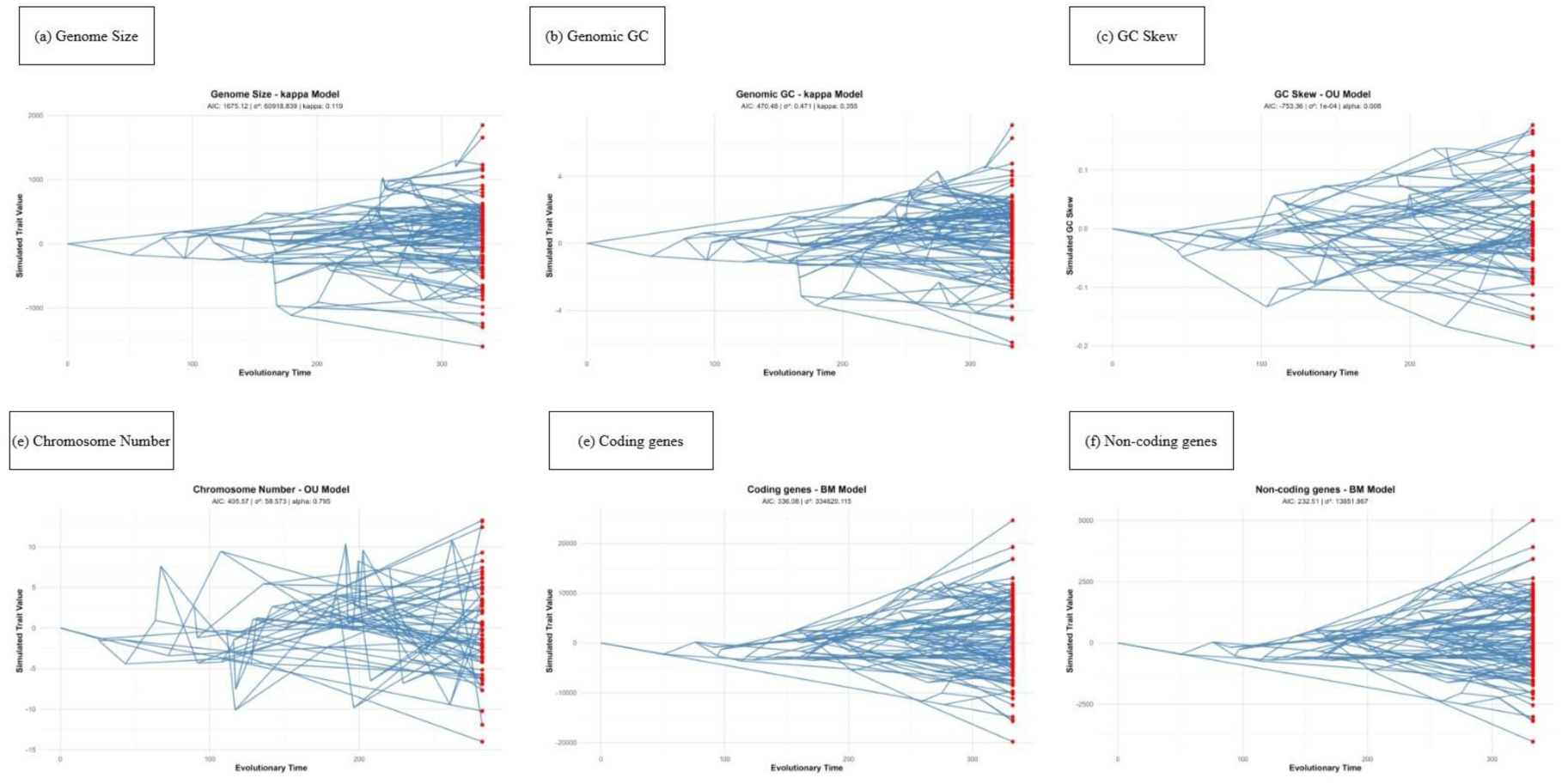
Simulated evolution of genomic traits in Coleoptera under best-fit evolutionary models.

Family-level genomic analyses revealed both congruence and divergence with order-level evolutionary patterns. While many families exhibited evolutionary modes consistent with their broader taxonomic orders, several lineages demonstrated distinctive trajectories (Supplementary Table 10 (ST10)). For instance, within *Lepidoptera*, genome size evolution followed constrained processes in *Hesperiidae* (OU, α=0.058, N_species=119) and *Nymphalidae* (OU, α=0.035, N_species=180), yet showed different evolutionary dynamics in *Lycaenidae* (kappa, κ=0.471, N_species=43). Notably, the low alpha values in OU models approach Brownian motion-like evolution, suggesting weaker constraint than initially apparent. Similarly, GC content conservation patterns varied in strength across families, with *Drosophilidae* (kappa, κ=0.686, N_species=235) showing particularly strong phylogenetic signal compared to other dipterans. This heterogeneity within orders underscores that lineage-specific factors can override broader taxonomic trends, with certain families exhibiting evolutionary modes distinct from their ordinal counterparts. In contrast, chromosome number evolution was best described by White Noise models across most families, consistent with the lack of phylogenetic structure expected for this trait. The prevalence of White Noise models for coding and non-coding gene counts in smaller datasets (*Aphididae* N_species=13, *Curculionidae* N_species=6) likely reflects inadequate sampling rather than true evolutionary patterns.

Our trait-specific analysis revealed a consistent pattern of increased kappa model prevalence in species-rich orders (*Coleoptera, Diptera, Hemiptera, Hymenoptera, Lepidoptera*), particularly for genome size (rich: 100% vs. poor: 17%) and Genomic GC (rich: 80% vs. poor: 0%). While statistical power was limited by sample sizes (Genomic GC: *P* = 0.071, Genome size: *P* = 0.107), the directional consistency across traits suggests that speciation-linked evolution may be more common in rapidly diversifying lineages. Species-poor orders (*Blattodea, Odonata, Orthoptera, Phasmatodea, Psocodea, Trichoptera*) showed greater reliance on gradual and adaptive radiation (BM and EB) models. Future studies with broader taxonomic sampling are needed to confirm these patterns statistically. Evolutionary model comparisons for genomic traits, including AIC values for all fitted models, are provided in Supplementary Table 11 (ST11).

### 3.3 PGLS -

#### A. Morphological Features

Phylogenetically corrected analyses (PGLS) revealed strong, order-specific patterns of morphological integration. Across all orders, structural traits (head dimensions, thorax width, body length) showed robust positive correlations, indicating deep evolutionary integration. For instance, head width and thorax width were tightly linked in *Coleoptera* (slope = 0.83, *P* < 0.001, R² = 0.89) and *Odonata* (slope = 1.02, *P* < 0.001, R² = 0.92).

In contrast, bite force demonstrated more variable and often weaker relationships. It was significantly correlated with other traits in some orders, such as *Orthoptera* (with wing length, *P* = 0.0003), but showed a unique negative correlation with thorax width in *Blattodea* (slope = - 33.93, *P* = 0.017). These results underscore that while a conserved pattern of structural integration exists, the functional relationship of bite force with body plan evolves in an order-specific manner. Phylogenetic generalized least squares (PGLS) results for morphological traits are presented in Supplementary Tables 12 (ST12) (significant relationships) and 13 (ST13) (all relationships).

#### B. Genomic features

Phylogenetically corrected analyses (PGLS) revealed that the relationship between core genomic features is highly order-specific. The correlation between genome size and GC content varied dramatically in both strength and direction across lineages. A significant negative correlation was detected in *Blattodea* (slope = −290.27, *P* = 0.038) and *Hymenoptera* (*P* = 0.006), while a strong positive correlation was found in *Odonata* (slope = 112.79, *P* = 0.012). Phasmatodea exhibited a significant negative relationship (slope = −349.12, *P* < 0.001, R² = 0.89).The slopes for GC skew versus genome size were essentially zero (*Lepidoptera*: slope ≈ 0, *P* < 0.001; *Hemiptera*: slope ≈ 0, *P* < 0.001), indicating statistically significant but minimal effect sizes, indicating that increasing genome size does not change the compositional asymmetry. Separately, chromosome number showed strong associations with gene content, exhibiting negative relationships in *Coleoptera* (slope = −24.61, *P* < 0.001) and positive relationships in *Phasmatodea* (slope = 12.80, *P* = 0.011).

These results underscore a lack of a universal evolutionary rule linking these fundamental genomic traits. Instead, the relationship between genome size and base composition appears to be governed by distinct, lineage-specific evolutionary pressures. Phylogenetic generalized least squares (PGLS) results for morphological traits are presented in Supplementary Tables 14 (ST14) (significant relationships) and 15 (ST15) (all relationships).

### 3.4 Importance of Morphological Features and Genomic Features in Determining Bite Force and Wing Length

Random Forest analysis identified key morphological predictors for bite force and wing length, revealing order-specific patterns. For bite force, wing length, and thorax width were consistently important (e.g., wing length importance = 9.57 in *Orthoptera*), suggesting a biomechanical link between flight musculature and biting power. Results of random forest analyses for bite force and wing length against other morphological traits are provided in Supplementary Tables 16 (ST16) and 17 (ST17), respectively.

However, phylogenetically-corrected linear models (*phylr*) revealed that evolutionary history strongly mediates these relationships. For bite force, most morphological traits exhibit no significant linear relationship across any of the orders, even after accounting for phylogeny (Supplementary Table 18 (ST18)). The few significant associations were order-specific, such as in *Odonata* (body length, *P* = 0.001). This indicates that the apparent morphological constraints on bite force are largely a consequence of shared evolutionary history rather than direct functional relationships. In contrast, for wing length, PhylR models showed strong, significant positive correlations with body size traits (e.g., body length, head height) across most orders (*P* < 0.001) ((Supplementary Table 19 (ST19))), which were corroborated by high importance scores in Random Forest analyses. This demonstrates that wing length evolves in a tightly integrated manner with overall body size, a pattern that remains robust even after correcting for phylogeny. Head parameters play a crucial role in influencing wing length, a pattern shown here in *Coleoptera* and *Hymenoptera* (Figure 8) and consistently recovered across other major insect lineages (10.5281/zenodo.17524136).

**Figure 8.**
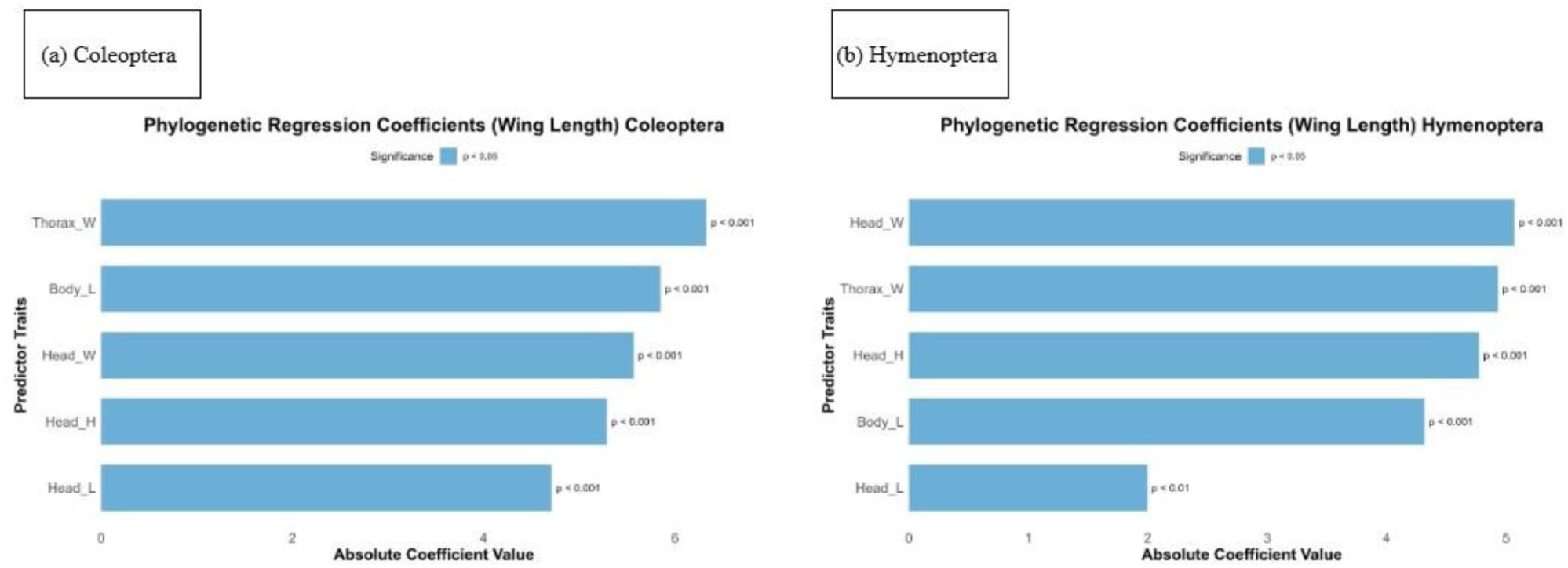
Phylogenetic mixed model results for wing length evolution. Conditional coefficients from phylogenetic regressions reveal the relative importance and effect sizes of morphological traits in predicting wing length for (A) Coleoptera and (B) Hymenoptera.

### 3.5 Testing for Pulsated Evolution

#### A. Morphological Traits

Kolmogorov-Smirnov and Anderson-Darling tests of phylogenetically independent contrasts show that pulsed evolution is a rare and order-specific phenomenon for morphological traits (Table 5). This pattern is underscored by the fact that only 16 out of a possible 70 trait-order combinations showed significant deviations.

**Table 5.**
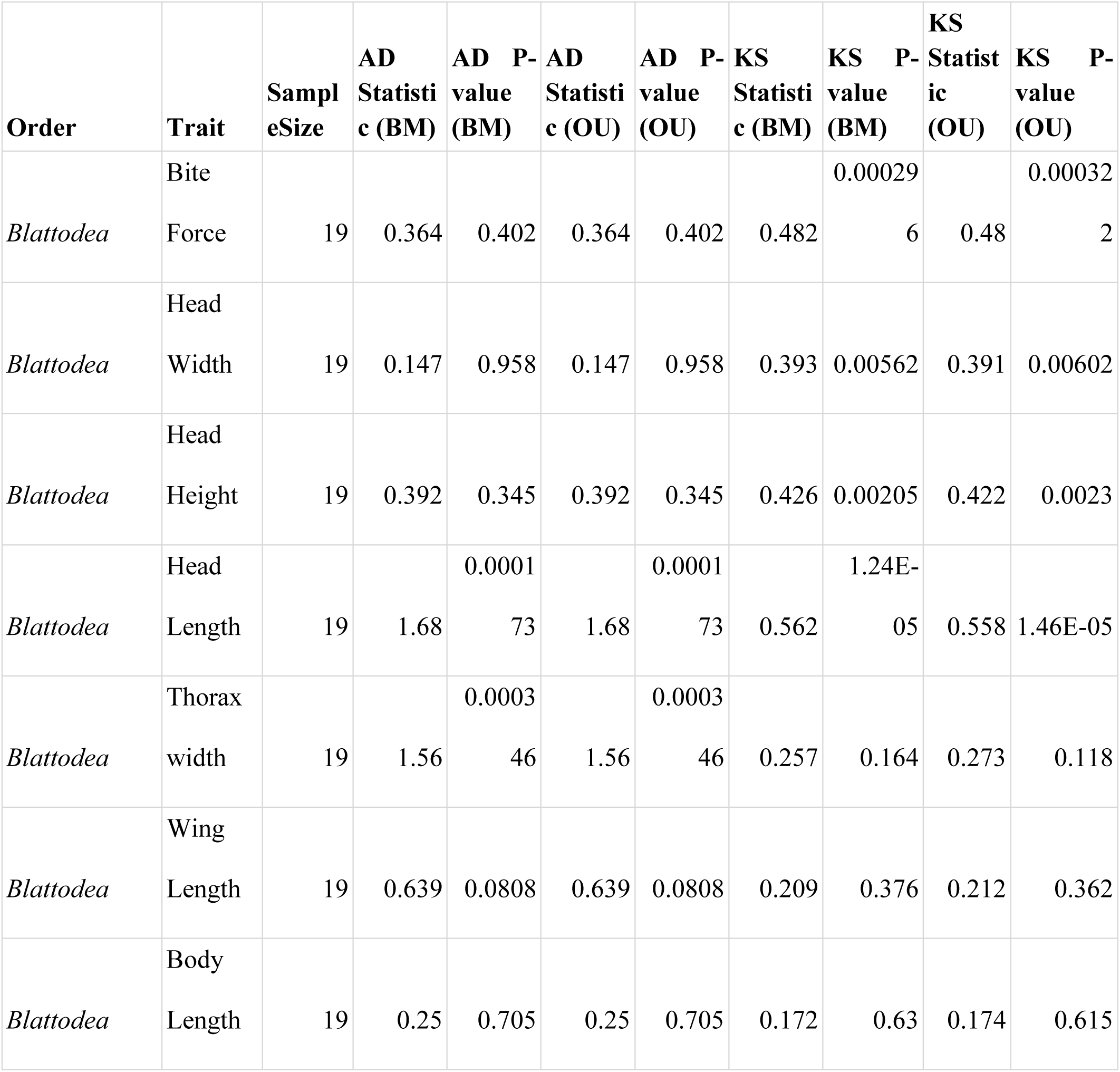
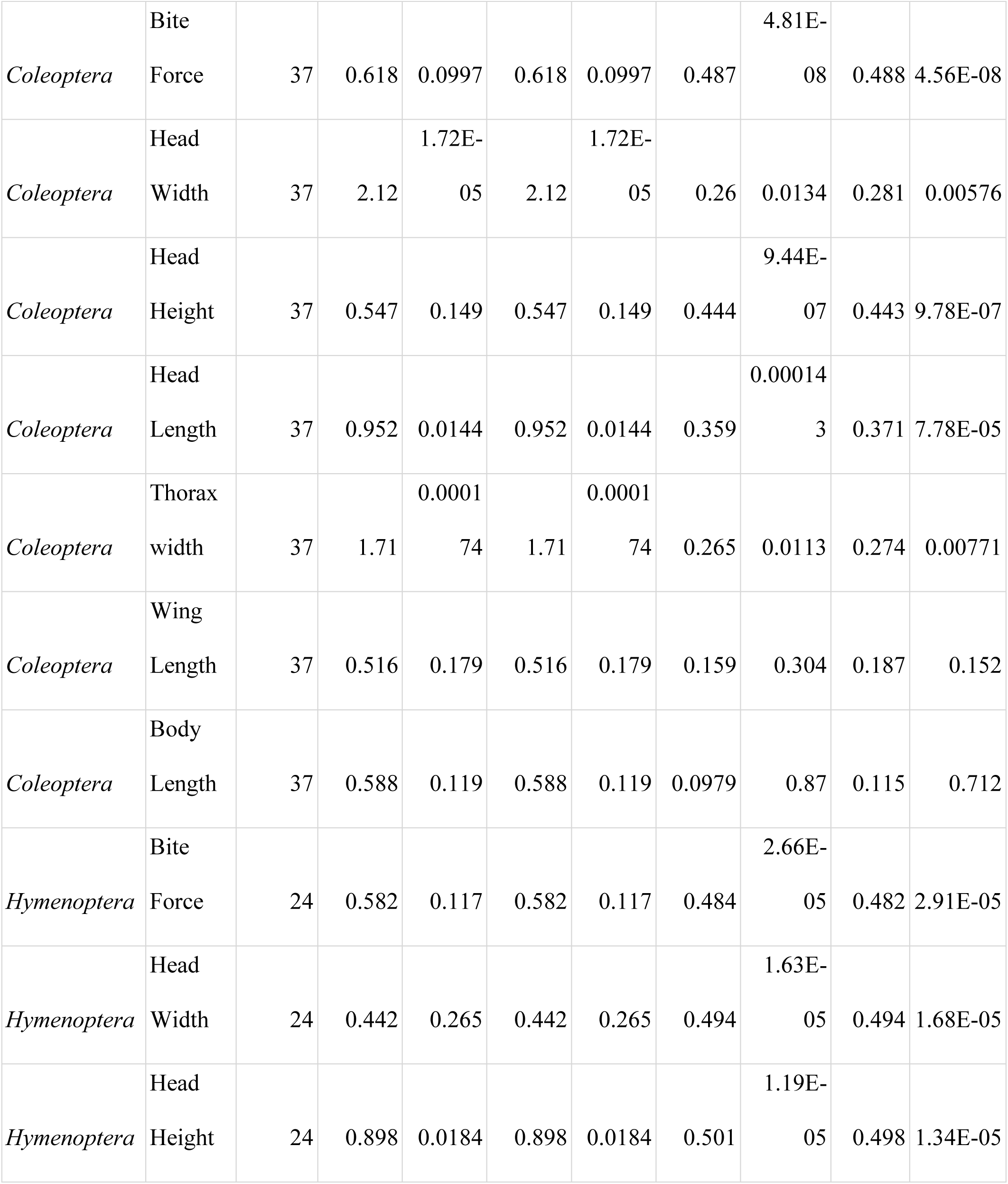
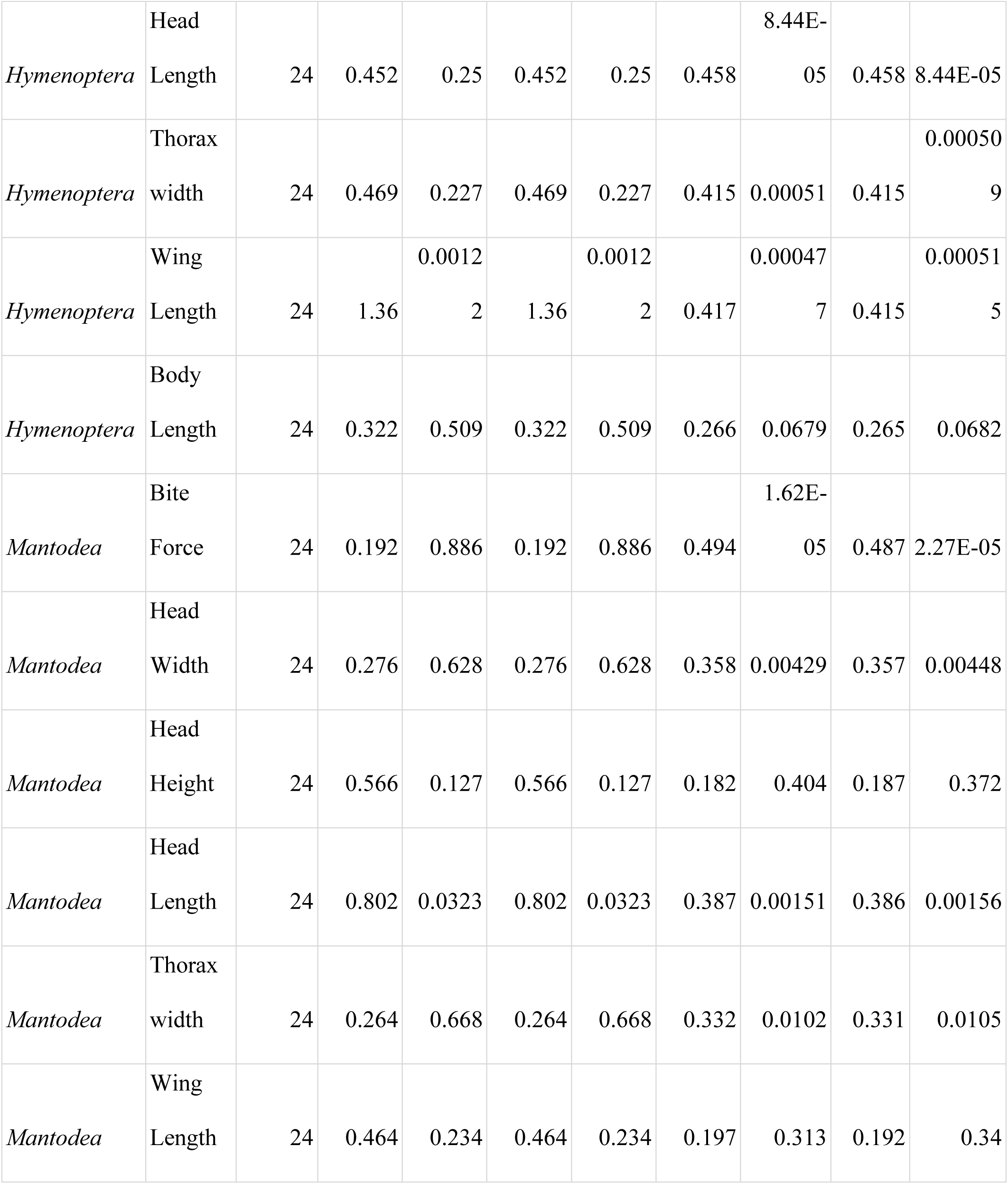
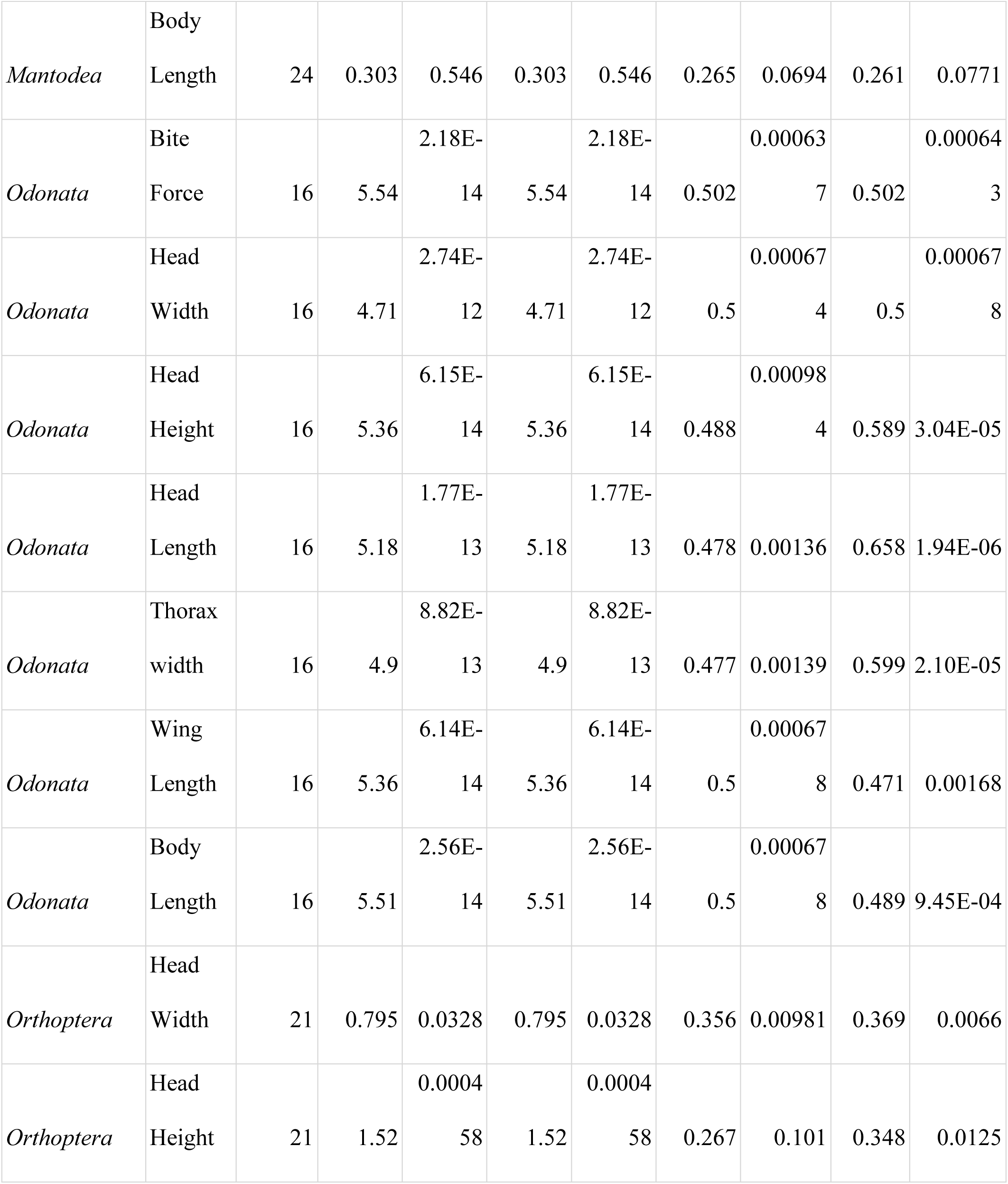
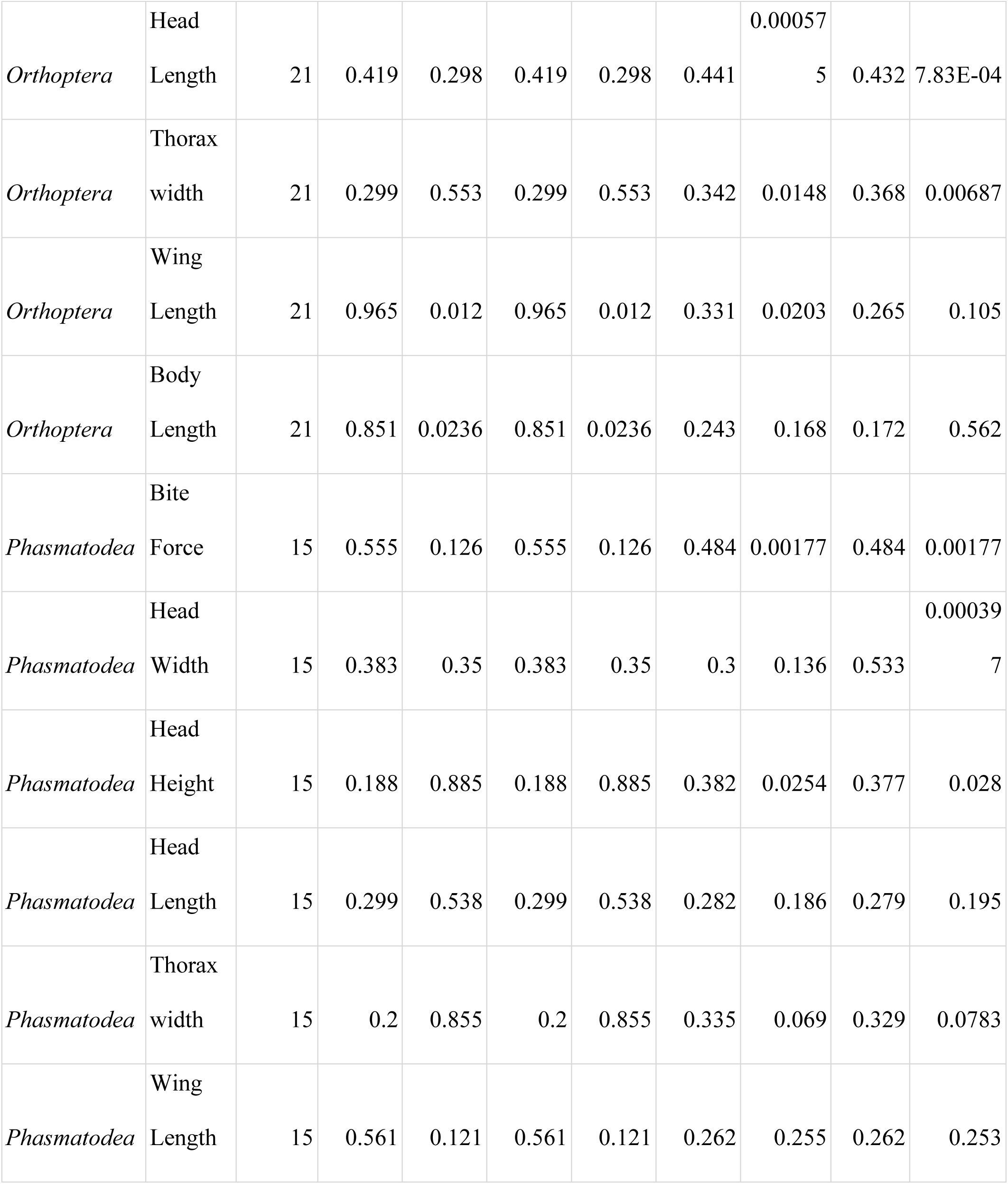
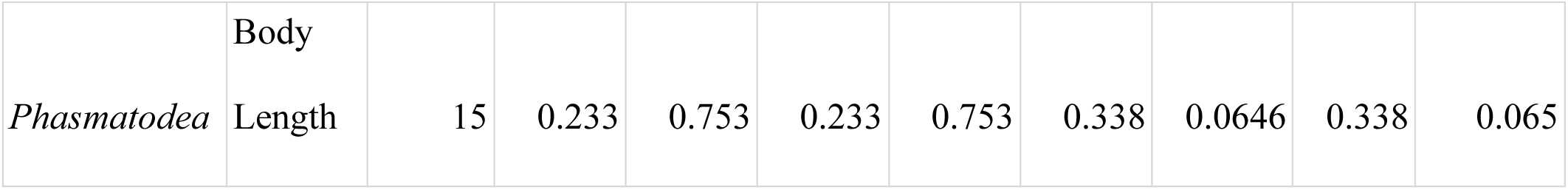
Statistical tests for pulsed evolution in morphological traits. Results of Anderson-Darling (AD) and Kolmogorov-Smirnov (KS) tests comparing the distribution of phylogenetically independent contrasts against the theoretical distributions of Brownian Motion (BM) and Ornstein-Uhlenbeck (OU) models

Most orders, including *Blattodea*, *Coleoptera,* and *Hymenoptera*, showed no significant deviations for the majority of their traits in both tests (all *P* > 0.05), indicating their evolution is well-described by gradual or stabilizing models. In contrast, *Mantodea* exhibited consistent evidence of pulsed evolution across both tests in head length (BM: KS = 0.284, *P* =0.033; AD = 0.802, *P* =0.032) and thorax width (BM: KS = 0.273, *P* = 0.046). The significant deviations from standard models in *Orthoptera* bite force across both tests correspond to its white noise evolutionary pattern, representing non-phylogenetic trait evolution rather than pulsed dynamics along lineages. Strikingly, *Odonata* was the only order where pulsed evolution was detected across all seven measured traits (Bite Force, Head Width, Head Height, Head Length, Thorax Width, Wing Length, and Body Length). The concentrated pulsed evolution in species-poor orders like *Mantodea* and *Odonata* supports a ’pulse mode of diversification’ (Ricklefs, 2013), where brief bursts of phenotypic and lineage diversification are followed by clade size decline. These results confirm that bursts of trait evolution are not a general rule but occur in specific lineages and for specific functional traits (Table 5). Figure 9 compares morphological trait evolution models for Coleoptera using phylogenetically independent contrasts, with complete PIC analyses for all orders available in Zenodo [10.5281/zenodo.17524136].

**Figure 9.**
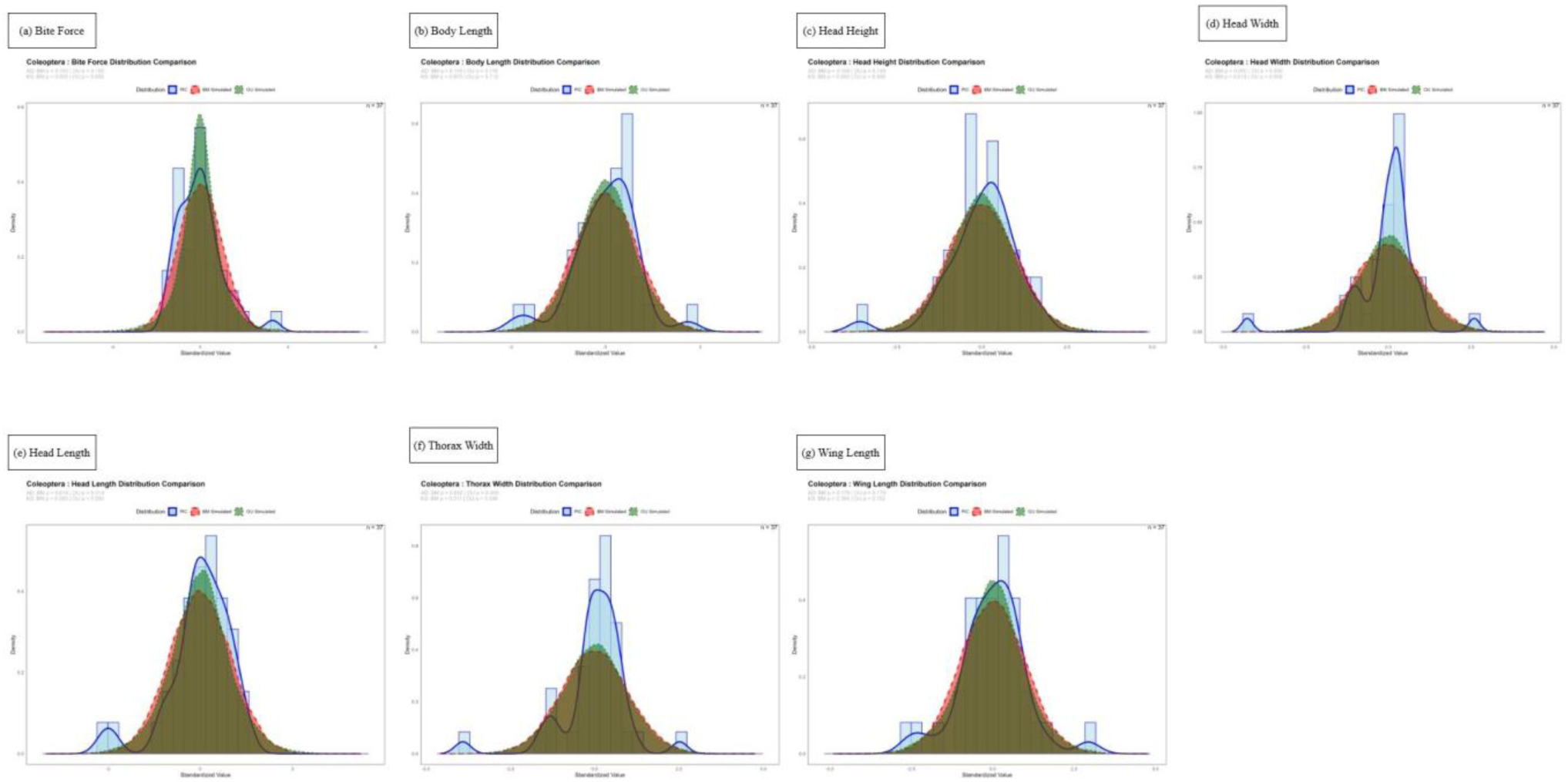
Model comparison of morphological trait evolution using phylogenetically independent contrasts (PICs). Using Coleoptera as a representative model, the distribution of standardized PICs is shown for (a) Bite Force, (b) Body Length, (c) Head Height, (d) Head Width, (e) Head Length, (f) Thorax Width, and (g) Wing Length, compared against expectations under Brownian motion (BM) and Ornstein-Uhlenbeck (OU) models.

#### B. Genomic Traits

Statistical analyses revealed that pulsed evolution is the dominant mode for genomic trait evolution. A significant majority of tests (60 out of 78, 77 %) rejected both Brownian Motion and Ornstein-Uhlenbeck models (p < 0.05) (Table 6). This pattern was pervasive across all well-sampled orders (n > 15), while the few non-significant results were confined to data-poor lineages (e.g., *Orthoptera,* n=7), indicating a lack of statistical power rather than an absence of the pattern. *Lepidoptera* showed the strongest signals, with all core genomic features deviating significantly from standard models (e.g., Chromosome Number: AD > 45, *P* < 3.7e-24). Similarly, *Coleoptera* and *Diptera* exhibited widespread pulses in traits like genome size (*Coleoptera* AD = 12.09; *Diptera* AD = 19.26; both *P* < 3.7E-24). Stronger signals from KS tests relative to AD tests in *Hemiptera* (e.g., Genome GC: KS *P* < 4.2E-22 vs. AD *P* = 0.0005) indicate that pulsed evolution in this order is characterized by systematic shifts in the distribution of evolutionary rates, rather than by the influence of extreme outliers. In contrast, orders like *Blattodea* and *Odonata* showed pulsed evolution only in specific features like GC skew.

**Table 6.** Statistical tests for pulsed evolution in genomic traits. Results of Anderson-Darling (AD) and Kolmogorov-Smirnov (KS) tests comparing the distribution of phylogenetically independent contrasts against the theoretical distributions of Brownian Motion (BM) and Ornstein-Uhlenbeck (OU) models

| Order | Trait | Sample Size | AD_St<br>atistic_<br>BM | AD_P_<br>Value_<br>BM | AD_St<br>atistic_<br>OU | AD_P_<br>Value_<br>OU | KS_Sta<br>tistic_B<br>M | KS_P_<br>Value_<br>BM | KS_Sta<br>tistic_<br>OU | KS_P_<br>Value_<br>OU |
| --- | --- | --- | --- | --- | --- | --- | --- | --- | --- | --- |
| *Blattodea* | Genome Size | 9 | 0.5351<br>2 | 0.1207<br>11 | 0.5351<br>2 | 0.1207<br>11 | 0.495 | 0.0246 | 0.488 | 0.0278 |
| *Blattodea* | Genomic GC | 9 | 0.3588<br>25 | 0.3650<br>68 | 0.3588<br>25 | 0.3650<br>68 | 0.555 | 0.0079<br>3 | 0.667 | 6.76E-04 |
| *Blattodea* | GC skew | 9 | 0.8087<br>28 | 0.0218<br>99 | 0.8087<br>28 | 0.0218<br>99 | 0.667 | 0.0006<br>76 | 0.667 | 6.76E-04 |
| *Coleoptera* | Genome Size | 109 | 12.086<br>33 | 3.70E-24 | 12.086<br>33 | 3.70E-24 | 0.238 | 9.05E-06 | 0.231 | 1.86E-05 |
| *Coleoptera* | Genomic GC | 109 | 5.8632<br>63 | 1.60E-14 | 5.8632<br>63 | 1.60E-14 | 0.193 | 0.0006<br>1 | 0.187 | 9.78E-04 |
| *Coleoptera* | GC skew | 75 | 13.377<br>45 | 3.70E-24 | 13.377<br>45 | 3.70E-24 | 0.338 | 7.43E-08 | 0.336 | 9.18E-08 |
| *Coleoptera* | Chromosome Number | 61 | 6.132281 | 2.97E-15 | 6.132281 | 2.97E-15 | 0.18 | 0.0387 | 0.138 | 0.199 |
| *Coleoptera* | Coding genes | 15 | 0.343662 | 0.438591 | 0.343662 | 0.438591 | 0.5 | 0.00113 | 0.502 | 1.05E-03 |
| *Coleoptera* | Non-coding genes | 12 | 0.221664 | 0.779914 | 0.221664 | 0.779914 | 0.496 | 0.00552 | 0.495 | 0.00556 |
| *Diptera* | Genome Size | 334 | 19.2557 | 3.70E-24 | 19.2557 | 3.70E-24 | 0.158 | 1.08E-07 | 0.155 | 2.40E-07 |
| *Diptera* | Genomic GC | 334 | 15.0611 | 3.70E-24 | 15.0611 | 3.70E-24 | 0.14 | 4.06E-06 | 0.138 | 5.92E-06 |
| *Diptera* | GC skew | 315 | 16.49241 | 3.70E-24 | 16.49241 | 3.70E-24 | 0.164 | 8.23E-08 | 0.162 | 1.44E-07 |
| *Diptera* | Chromosome Number | 106 | 13.56495 | 3.70E-24 | 13.56495 | 3.70E-24 | 0.235 | 1.59E-05 | 0.302 | 8.23E-09 |
| *Diptera* | Coding genes | 76 | 3.100458 | 7.50E-08 | 3.100458 | 7.50E-08 | 0.5 | 6.32E-17 | 0.498 | 8.52E-17 |
| *Diptera* | Non - coding genes | 74 | 8.3568<br>11 | 1.58E-20 | 8.3568<br>11 | 1.58E-20 | 0.498 | 2.34E-16 | 0.5 | 1.89E-16 |
| *Hemiptera* | Genome Size | 74 | 21.284<br>7 | 3.70E-24 | 21.284<br>7 | 3.70E-24 | 0.493 | 4.66E-16 | 0.486 | 1.32E-15 |
| *Hemiptera* | Genomic GC | 74 | 1.5602<br>13 | 4.71E-04 | 1.5602<br>13 | 4.71E-04 | 0.315 | 8.15E-07 | 0.581 | 4.17E-22 |
| *Hemiptera* | GC skew | 41 | 1.4317<br>29 | 9.11E-04 | 1.4317<br>29 | 9.11E-04 | 0.16 | 0.243 | 0.177 | 0.156 |
| *Hemiptera* | Coding genes | 23 | 1.5055<br>34 | 5.14E-04 | 1.5055<br>34 | 5.14E-04 | 0.499 | 2.16E-05 | 0.5 | 2.08E-05 |
| *Hemiptera* | Non - coding genes | 20 | 1.0194<br>43 | 0.0085<br>98 | 1.0194<br>43 | 0.0085<br>98 | 0.498 | 9.80E-05 | 0.493 | 1.21E-04 |
| *Hemiptera* | GC_skew | 68 | 5.3068<br>03 | 3.06E-13 | 5.3068<br>03 | 3.06E-13 | 0.691 | 1.30E-28 | 0.691 | 1.30E-28 |
| *Hymenoptera* | Genome Size | 169 | 16.115<br>95 | 3.70E-24 | 16.115<br>95 | 3.70E-24 | 0.228 | 4.74E-08 | 0.226 | 6.16E-08 |
| *Hymenoptera* | Genomic GC | 169 | 1.7329<br>49 | 1.86E-04 | 1.7329<br>49 | 1.86E-04 | 0.0827 | 0.198 | 0.0795 | 0.236 |
| *Hymenoptera* | GC skew | 157 | 13.45811 | 3.70E-24 | 13.45811 | 3.70E-24 | 0.209 | 2.35E-06 | 0.204 | 4.21E-06 |
| *Lepidoptera* | Genome Size | 585 | 14.50234 | 3.70E-24 | 14.50234 | 3.70E-24 | 0.116 | 2.79E-07 | 0.115 | 3.67E-07 |
| *Lepidoptera* | Genomic GC | 585 | 11.77355 | 3.70E-24 | 11.77355 | 3.70E-24 | 0.103 | 9.28E-06 | 0.102 | 1.18E-05 |
| *Lepidoptera* | GC skew | 565 | 23.72859 | 3.70E-24 | 23.72859 | 3.70E-24 | 0.142 | 2.33E-10 | 0.142 | 2.39E-10 |
| *Lepidoptera* | Chromosome Number | 311 | 45.46504 | 3.70E-24 | 45.46504 | 3.70E-24 | 0.276 | 6.57E-21 | 0.276 | 5.43E-21 |
| *Lepidoptera* | Coding_genes | 42 | 5.056644 | 9.98E-13 | 5.056644 | 9.98E-13 | 0.29 | 0.00171 | 0.288 | 1.86E-03 |
| *Lepidoptera* | Non_coding_genes | 37 | 1.076688 | 0.006976 | 1.076688 | 0.006976 | 0.158 | 0.315 | 0.158 | 0.316 |
| *Odonata* | Genome Size | 11 | 0.173622 | 0.901488 | 0.173622 | 0.901488 | 0.473 | 0.0147 | 0.477 | 0.0136 |
| *Odonata* | Genomic GC | 11 | 0.411601 | 0.280129 | 0.411601 | 0.280129 | 0.631 | 0.000317 | 0.636 | 2.73E-04 |
| *Odonata* | GC skew | 11 | 1.21138 | 0.00205 | 1.21138 | 0.00205 | 0.727 | 1.79E-05 | 0.727 | 1.79E-05 |
| *Orthoptera* | Genome Size | 7 | - | - | - | - | 0.246 | 0.792 | 0.245 | 0.795 |
| *Orthoptera* | Genome GC | 7 | - | - | - | - | 0.257 | 0.744 | 0.261 | 0.726 |
| *Phasmatodea* | Genome Size | 11 | 1.32969 | 9.90E-04 | 1.32969 | 9.90E-04 | 0.5 | 0.00817 | 0.502 | 7.83E-03 |
| *Phasmatodea* | Genome GC | 11 | 1.195104 | 0.002266 | 1.195104 | 0.002266 | 0.354 | 0.128 | 0.495 | 9.1E-03 |

The overwhelming prevalence of pulsed evolution in genomic architecture, consistently supported by both AD and KS tests, suggests that large-scale genomic changes represent a fundamental mechanism of lineage diversification in insects, often superseding gradual models of evolution (Table 6). Figure 10 compares genomic trait evolution models for Coleoptera using phylogenetically independent contrasts, while PIC analyses for all other orders are archived in Zenodo [10.5281/zenodo.17524136].

**Figure 10.**
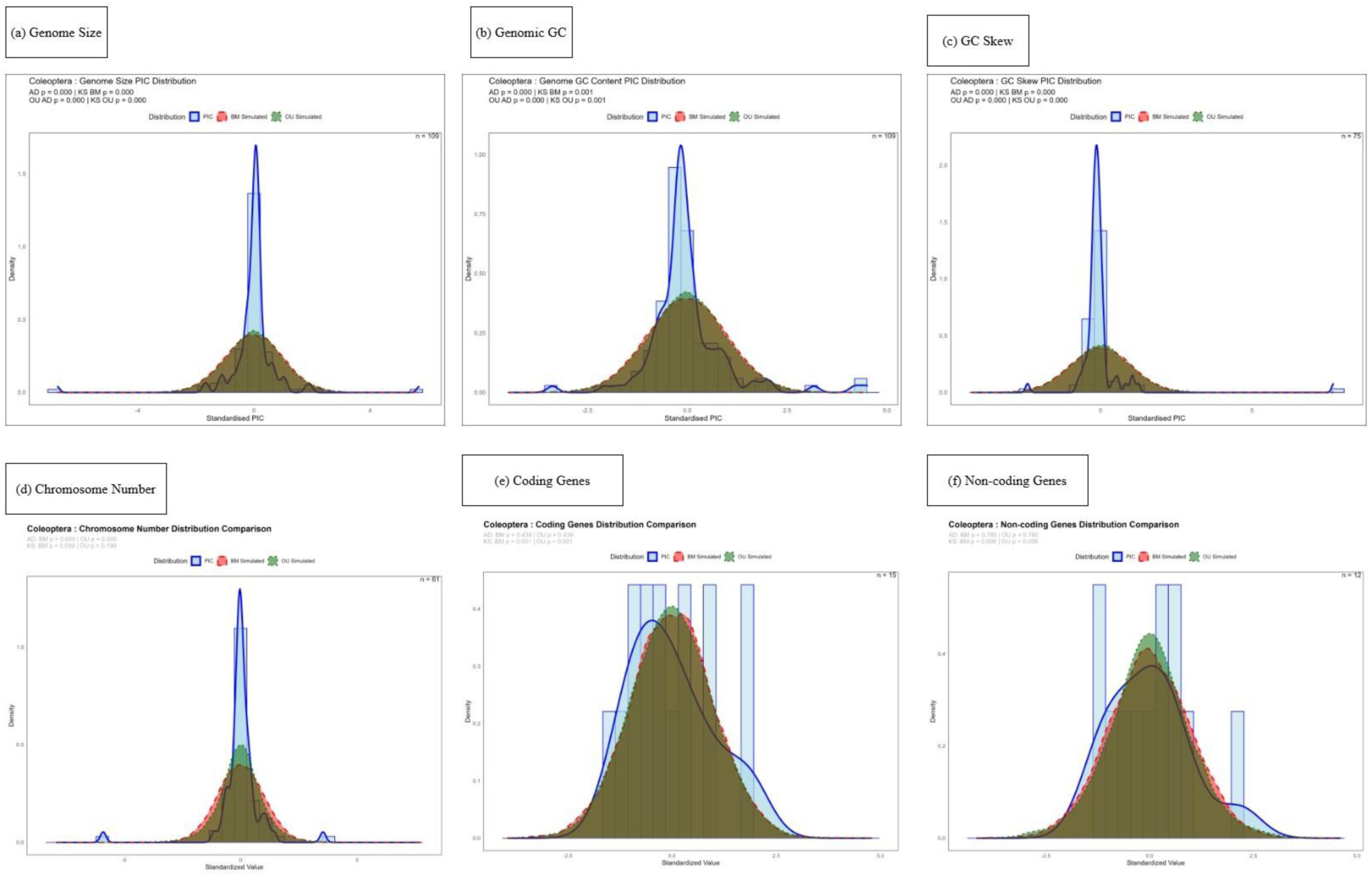
Model comparison of genomic trait evolution using phylogenetically independent contrasts (PICs). Using Coleoptera as a representative model, the distribution of standardized PICs is shown for (a) Genome Size, (b) Genomic GC, (c) GC Skew, (d) Chromosome Number, (e) Coding Genes, and (f) Non-coding Genes, compared against expectations under Brownian motion (BM) and Ornstein-Uhlenbeck (OU) models.

### Pulses in Morphological Trait Changes Across Insect Orders

The patchwork distribution of significant pulsed evolution signals (ST20) highlights its lineage-specific nature. A re-evaluation of model support (Table 7) clarifies these patterns: head length evolution is pulsed in *Coleoptera* (ΔAIC = −31.4) and *Mantodea* (ΔAIC = −46.2), but not gradual in the latter as initially suggested. Similarly, thorax width evolution is best fit by a pulsed model in *Coleoptera* (ΔAIC = −33.7) and *Odonata* (ΔAIC = −79.8), while wing length follows a pulsed model in *Odonata* (ΔAIC = −45.4) but not in *Hymenoptera*, where Brownian motion is preferred (ΔAIC = +17.8). This contingent model-fitting underscores that pulses are not trait-autonomous but depend on clade-specific selective regimes or historical contingencies. The exception is *Odonata*, where pulsed models were selected for six of seven traits, indicating a unique, order-wide dynamic. Further supporting this mosaic pattern, *Orthoptera* shows pulsed evolution for head length (ΔAIC = −15.8) and head width, while Brownian motion characterizes the evolution of its bite force and body length.

**Table 7.**
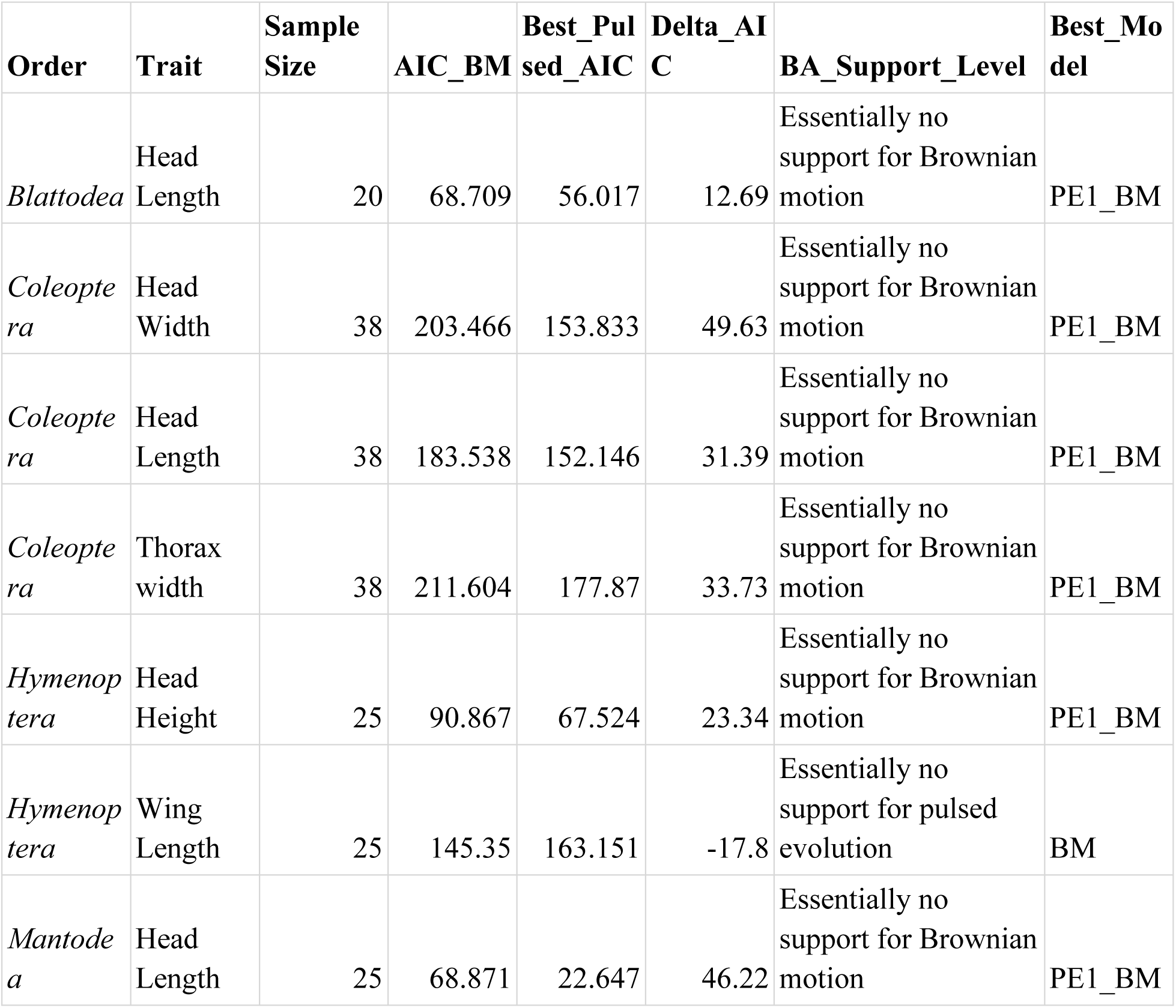
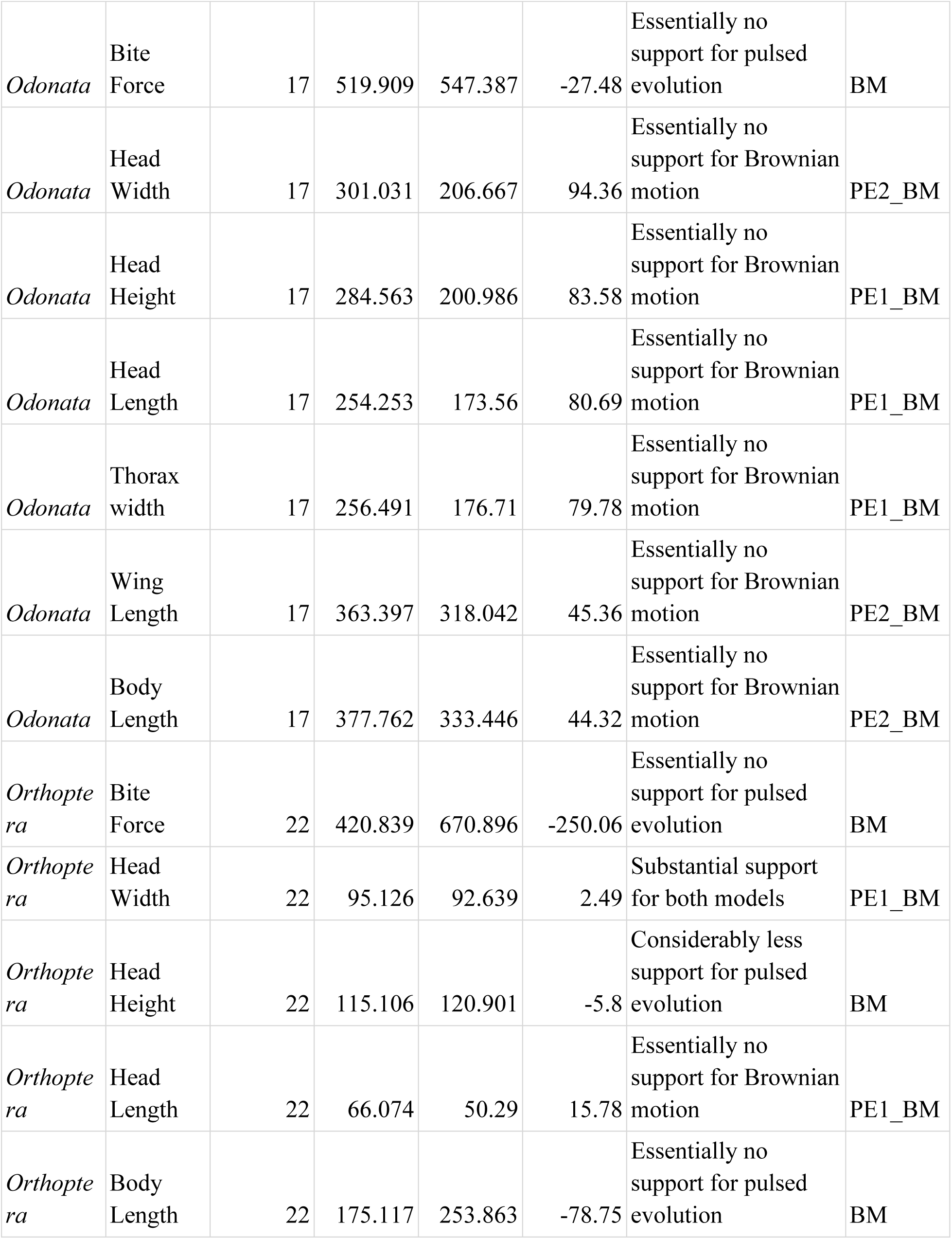
Model selection results for morphological trait evolution. Models compare a Brownian motion (BM) process to pulsed evolution (PE) models. The best-fitting model for each trait, based on Akaike Information Criterion (AIC) scores, is indicated.

| Order | Trait | Sample Size | AIC_BM | Best_Pulsed_AIC | Delta_AIC | BA_Support_Level | Best_Model |
| --- | --- | --- | --- | --- | --- | --- | --- |
| *Blattodea* | Head Length | 20 | 68.709 | 56.017 | 12.69 | Essentially no support for Brownian motion | PE1_BM |
| *Coleoptera* | Head Width | 38 | 203.466 | 153.833 | 49.63 | Essentially no support for Brownian motion | PE1_BM |
| *Coleoptera* | Head Length | 38 | 183.538 | 152.146 | 31.39 | Essentially no support for Brownian motion | PE1_BM |
| *Coleoptera* | Thorax width | 38 | 211.604 | 177.87 | 33.73 | Essentially no support for Brownian motion | PE1_BM |
| *Hymenoptera* | Head Height | 25 | 90.867 | 67.524 | 23.34 | Essentially no support for Brownian motion | PE1_BM |
| *Hymenoptera* | Wing Length | 25 | 145.35 | 163.151 | -17.8 | Essentially no support for pulsed evolution | BM |
| *Mantodea* | Head Length | 25 | 68.871 | 22.647 | 46.22 | Essentially no support for Brownian motion | PE1_BM |
| *Odonata* | Bite Force | 17 | 519.909 | 547.387 | -27.48 | Essentially no support for pulsed evolution | BM |
| *Odonata* | Head Width | 17 | 301.031 | 206.667 | 94.36 | Essentially no support for Brownian motion | PE2_BM |
| *Odonata* | Head Height | 17 | 284.563 | 200.986 | 83.58 | Essentially no support for Brownian motion | PE1_BM |
| *Odonata* | Head Length | 17 | 254.253 | 173.56 | 80.69 | Essentially no support for Brownian motion | PE1_BM |
| *Odonata* | Thorax width | 17 | 256.491 | 176.71 | 79.78 | Essentially no support for Brownian motion | PE1_BM |
| *Odonata* | Wing Length | 17 | 363.397 | 318.042 | 45.36 | Essentially no support for Brownian motion | PE2_BM |
| *Odonata* | Body Length | 17 | 377.762 | 333.446 | 44.32 | Essentially no support for Brownian motion | PE2_BM |
| *Orthoptera* | Bite Force | 22 | 420.839 | 670.896 | -250.06 | Essentially no support for pulsed evolution | BM |
| *Orthoptera* | Head Width | 22 | 95.126 | 92.639 | 2.49 | Substantial support for both models | PE1_BM |
| *Orthoptera* | Head Height | 22 | 115.106 | 120.901 | -5.8 | Considerably less support for pulsed evolution | BM |
| *Orthoptera* | Head Length | 22 | 66.074 | 50.29 | 15.78 | Essentially no support for Brownian motion | PE1_BM |
| *Orthoptera* | Body Length | 22 | 175.117 | 253.863 | -78.75 | Essentially no support for pulsed evolution | BM |

The parameter estimates for morphological traits under pulsed evolutionary models reveal a fundamental divergence in dynamics. The order *Odonata*, despite its comparatively limited species diversity, is characterized by a variable-magnitude, multi-pulse regime (PE2_BM models featuring distinct σ1 and σ2 parameters). This contrasts with the simpler, low-rate, single-pulse pattern (PE1_BM) recovered for the remaining orders, which experienced more uniform pulsed changes throughout their evolutionary history. Supplementary Table 20 (ST20) catalogs the complete parameter set for the best-fitting pulsed evolution models of genomic traits across insect orders

### Pulses in Genomic Trait Changes Across Insect Orders

Our initial assessment of trait distributions (Table 6) using Anderson-Darling and Kolmogorov-Smirnov tests revealed widespread significant deviations from Brownian Motion and Ornstein-Uhlenbeck expectations for most genomic traits. However, a critical examination of evolutionary mode using Akaike Information Criterion (AIC) in Table 8 and Supplementary Table 21 (ST21) demonstrates that this widespread non-normality does not universally translate to support for pulsed evolution. For instance, while genome size in *Lepidoptera, Hemiptera, Odonata*, and *Phasmatodea* strongly deviated from normality (Table 6), the AIC provides decisive support for Brownian Motion as the best model, not a pulsed one. Conversely, the AIC reveals extremely strong, unambiguous support for pulsed evolution in traits like GC skew across nearly all orders (e.g., *Coleoptera, Diptera, Hymenoptera, Lepidoptera*) and for genome size itself in *Coleoptera, Diptera*, and *Hymenoptera*—cases where the initial normality tests were also significant. This AIC-based re-evaluation clarifies that the dominant evolutionary mode is highly trait- and lineage-specific: pulsed models powerfully explain variation in base composition (GC, GC skew) and architecture (chromosome number), Brownian motion is consistently the best fit for gene content, and genome size evolution is split, following a pulsed model in some major orders (especially the species rich ones) and a gradual one in others.

**Table 8.**
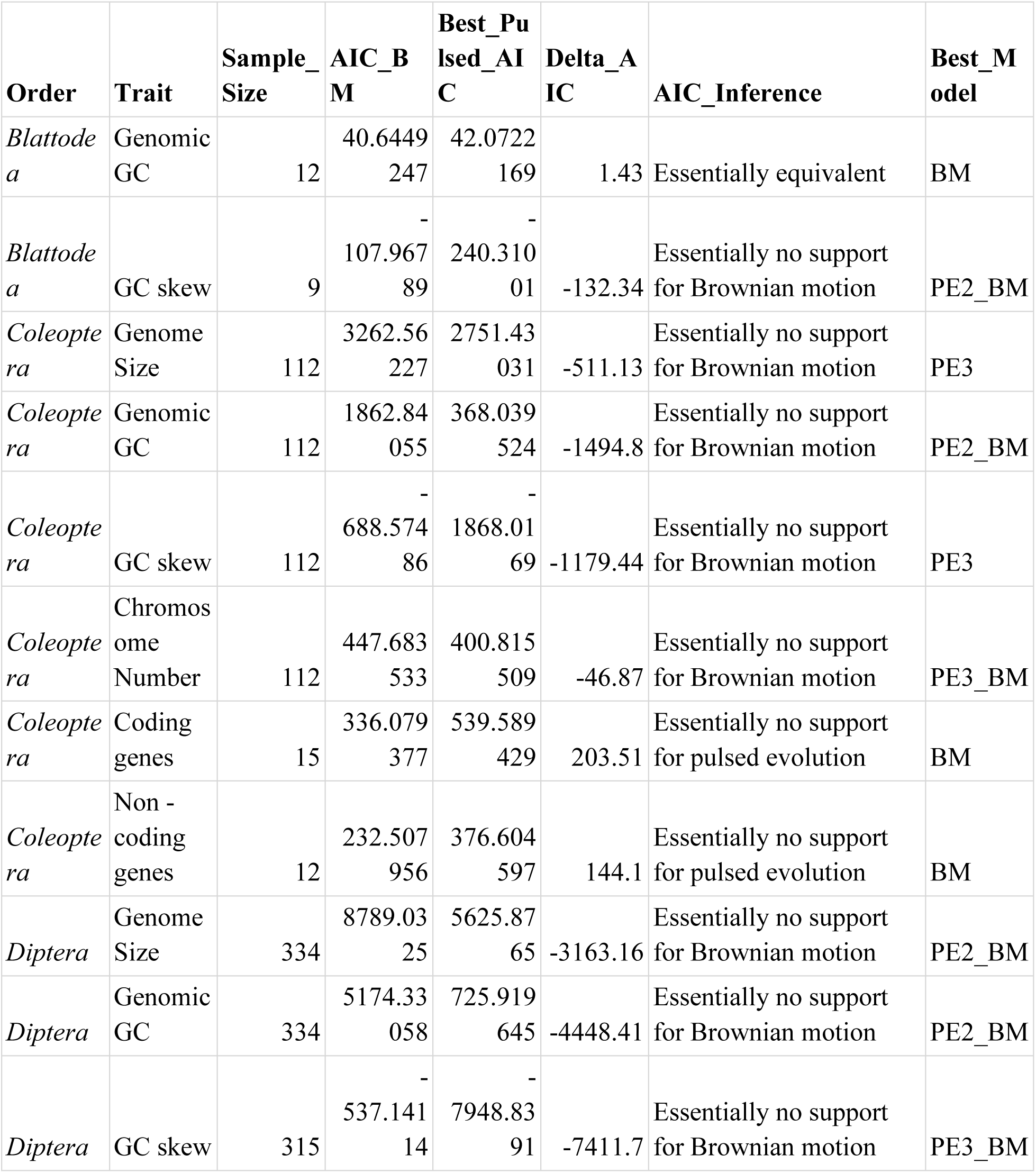
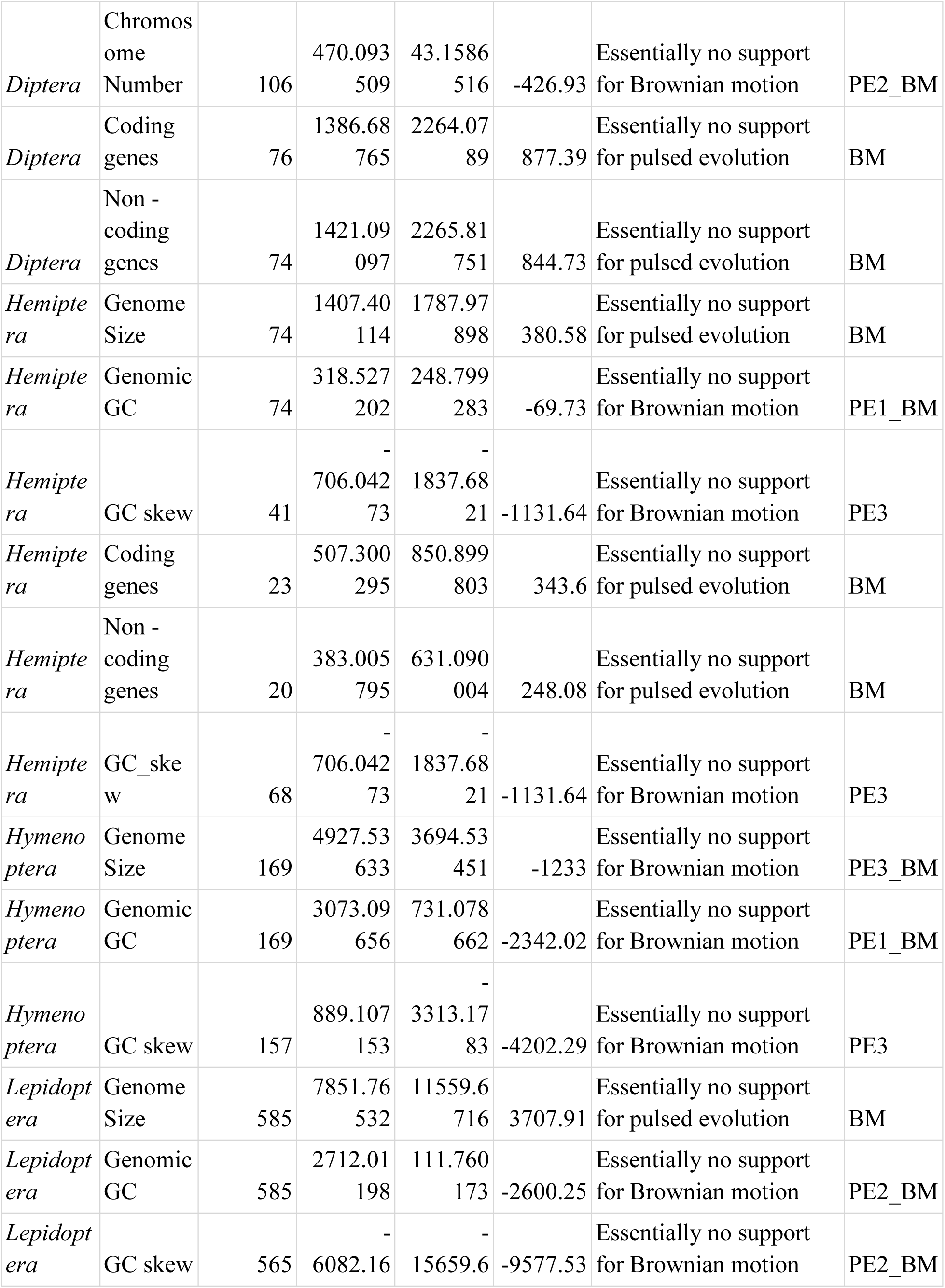
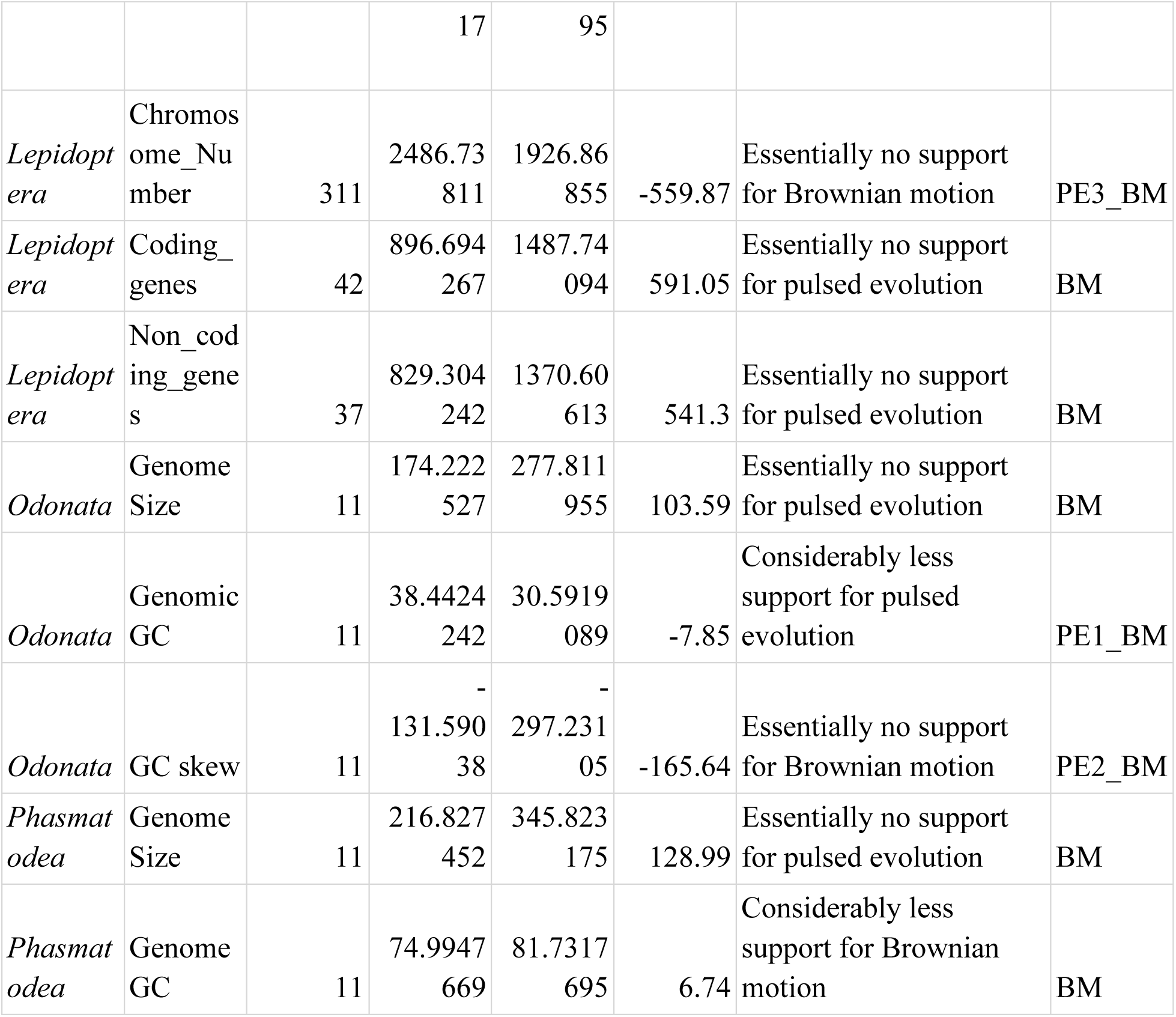
Model selection results for genomic trait evolution. Models compare a Brownian motion (BM) process to pulsed evolution (PE) models. The best-fitting model for each trait, based on Akaike Information Criterion (AIC) scores, is indicated.

The pattern of pulses in genomic trait parameters reveal a macroevolutionary dynamic that contrasts sharply with the morphological pattern. Here, pulsed evolution is pervasive across both species-rich and species-poor orders, with complex, multi-rate models (PE2_BM, PE3, PE3_BM) being the rule rather than the exception. Notably, species-rich orders frequently exhibit the most substantial genomic shifts, with Diptera and Hymenoptera showing exceptionally large σ values for genome size, indicative of profound, punctuated changes in genomic architecture. Unlike the morphology, where a simple, low-rate pulsed model characterized most orders, the prevalence of complex pulsed modes in genomic evolution across the insect tree of life suggests these processes may interact with lineage diversification in ways that are distinct from morphological evolution. Supplementary Table 21 (ST21) catalogs the complete parameter set for the best-fitting pulsed evolution models of genomic traits across insect orders

## Discussion

Our analyses across insect orders and families reveal important patterns of trait evolution, yet these findings must be interpreted within the context of data sampling limitations. The availability of species in time-calibrated phylogenies, coupled with uneven representation of morphological and genomic traits in public datasets, has resulted in heterogeneous sampling across the insect tree of life. While certain orders and traits exhibit adequate coverage for robust phylogenetic inference, others suffer from limited taxonomic sampling that may constrain the generalizability of our results. While broader taxonomic scales may obscure trait adaptations significant at finer ecological levels (Edel et al., 2024), our family-level analyses—though limited by sampling constraints—sought to uncover these nuanced evolutionary patterns.

Body mass-normalized bite force represents a fundamental metric in functional ecology (Christainsen & Wroe, 2007; Isip et al., 2022), reflecting dietary adaptations across species (Christainsen & Wroe, 2007; Christainsen, 2007; Santana et al., 2010; Tan et al., 2021). Our analyses demonstrate that even absolute bite force exhibits significant phylogenetic lability, suggesting its important role in dietary niche diversification (Edel et al.,2024), defense mechanisms (Buezas et al., 2019; Ruhr et al., 2024; Saif et al., 2025), and digging behaviors (Buezas et al., 2019), though we acknowledge potential confounding effects from body size (Isip et al., 2022), mandibular strength (Buezas et al., 2019), and other unmeasured morphological traits. This pattern of trait evolution aligns with phylogenetic niche conservatism (Losos, 2008; Wiens, 2011), where labile morphological traits enable ecological divergence into adjacent niches while maintaining broader evolutionary constraints on other organismal features. Consistent with Edel et al. (2024), who found no phylogenetic signal in mandibular action potential (MAP) in *Orthoptera*, our study similarly reveals an absence of phylogenetic signal in bite force across *Orthoptera* and most other insect orders examined. The persistence of a strong phylogenetic signal in bite force within the ecologically hyper-diverse *Hymenoptera* is a key finding, suggesting deep evolutionary constraints (Santos-Pinto et al., 2018). The same pattern in *Blattodea* aligns with expectations for a lineage with lower diversification and a more conserved niche (Ackerly, 2009). The evolution of bite force in *Hymenoptera* and *Blattodea* follows a Brownian Motion (BM) model without evidence of pulsed evolution, indicating a process dominated by either neutral drift or strong phylogenetic constraints. Bite force evolution correlates with thorax width in *Blattodea,* suggesting body plan constraints, but shows no relationship with other phylogenetically morphological traits in the remaining insect orders, indicating low phylogenetic conservatism.

Head length evolution is highly order-specific, forming a continuum from strong, pulsed phylogenetic structure in Coleoptera and Mantodea to weak or non-existent signal in Hymenoptera, Blattodea, Orthoptera, and Odonata. Interestingly, PGLS results show significant evolutionary correlations with other cranial traits, revealing coordinated evolution within a cranial module despite the lack of phylogenetic signal in head length itself. This pattern of modular integration, where traits evolve in concert, parallels observations in mammalian or amphibian, or reptilian crania (Porto et al., 2009; Bardua et al., 2019; Felis et al., 2019) and aligns with the developmental control of overall body size in Insects (Nijhout et al., 2014). All other measured morphological traits—head width, head height, thorax width, wing length, and body length— exhibit strong phylogenetic signals across the insect orders studied.

Wing length exhibits a significant phylogenetic signal across most datasets, indicating evolutionary trait conservation. This pattern is consistent with Ospina-Garcés et al., 2018, which found strong phylogenetic signal for Wing Length in *Scarabaeinae* but no correlating variables, whereas we find correlation with head and thorax traits. The absence of signal in the Staton et al. (2023) dataset is likely an artifact of sampling or experimental bias. The trait’s best-fitting evolutionary model varies, encompassing Brownian Motion (BM), Early Burst (EB), and Delta models, which implies divergent evolutionary processes across clades. Comparative analyses (PGLS, Phylogenetic mixed models) confirm that wing length is tightly integrated with the rest of the morphological traits, a pattern distinct from the decoupled evolution of bite force. Our analysis, leveraging phylogenetically-informed models, delineates that head parameters are the strongest morphological predictors of wing length. This finding of tight integration reflects the deeply conserved developmental coordination of the insect body plan (Oliveria et al., 2014; Shatkovska & Ghazali, 2016), illustrating the constrained allometric relationships that govern its architecture (Nijhout & Cavalier, 2015). By quantitatively establishing the primacy of head dimensions, our study adds a crucial morphological specificity to this established evolutionary-developmental principle.

Our distributional analyses reveal that pulsed evolution is infrequent and trait-specific rather than a widespread phenomenon in insect morphological evolution. This finding is consistent with Harmon et al. (2010), who similarly observed that punctuated diversification of morphological traits is rare across animal clades. The concentrated pulsed evolution in species-poor orders like Mantodea and Odonata supports a ’pulse mode of diversification’ (Ricklefs, 2013), where bursts of innovation are followed by clade size decline. This pattern aligns more with ecological control hypotheses, which predict a negative diversity-dependence, than with ’diversity begets diversity’ models (Whittaker et al., 2007; Ricklefs, 2013; Madi et al, 2020). While most traits across orders conform to Brownian motion or Ornstein-Uhlenbeck expectations, a limited subset—including head dimensions in *Coleoptera* and *Blattodea*, and wing length in *Hymenoptera* and *Mantodea* — show significant deviations consistent with evolutionary pulses. These findings suggest that certain functional traits, particularly those related to sensory and locomotor systems, may be more prone to episodic bursts of change that contribute disproportionately to morphological diversity. Future work should examine how these trait pulses correlate with environmental changes and enable niche partitioning. However, the overall pattern emphasizes developmental and functional constraints that maintain evolutionary stability in most morphological systems over deep timescales. The strong phylogenetic signal we observed in core morphological traits aligns with broader patterns in insect evolution, where even complex structures like genitalia and behaviors have been shown to exhibit significant phylogenetic conservatism (Song & Bucheli, 2010; Mayhew, 2018). For morphological traits, neither the strength of phylogenetic signal nor the specific mode of evolution emerges as a strong determinant of species richness. Thus, the macroevolutionary processes controlling trait evolution appear largely disconnected from those governing lineage diversification.

The evolutionary mode of genome size across insect orders is highly contingent rather than uniform. While a strong phylogenetic signal is evident, model selection demonstrates that the underlying process is split between pulsed and gradual evolution in a lineage-specific manner. Crucially, this split reveals a major macroevolutionary divide: the most species-rich and ecologically dominant orders—*Coleoptera*, *Diptera*, and *Hymenoptera*—consistently show strong support for pulsed evolution of their genome sizes. This pattern in such prolific clades aligns with dynamics observed in rapidly diversifying microbial lineages (Gao & Wu, 2022), suggesting pulsed evolution may be a significant mechanism linked to heightened diversification. However, the equally strong support for Brownian motion in other orders like Lepidoptera and Hemiptera indicates that pulsed evolution, while prominent in major radiations, is not the universal process governing genome size across all insects. Furthermore, the pulsed evolutionary mode is even more pronounced and widespread for genomic GC content than for genome size. Unlike genome size, where pulsed evolution is largely restricted to species-rich orders, pulsed dynamics in GC content are recovered across both species-rich and species-poor clades. The significant correlation between these two fundamental genomic traits, which persists after phylogenetic correction, underscores their coordinated evolution, a pattern observed across the tree of life (Alfsnes et al., 2017; Yuan et al., 2021). Crucially, our discovery that GC content is pervasively governed by pulsed evolution in insects provides a dynamic mechanistic explanation for this correlation and strongly aligns with the macroevolutionary pattern recently described in microbial lineages by Gao & Wu (2022). However, the macroevolutionary relationship between these traits is not uniform; it varies significantly across lineages. We found positive evolutionary correlations in Lepidoptera and Odonata, but negative correlations in Blattodea and Phasmatodea, revealing distinct, order-specific genomic evolutionary regimes underlying the broader trend of coordinated evolution. While previous studies have established the link between genome size and repetitive elements like transposons in insects (Yuan et al., 2021; Heckenhauer et al., 2022; Salman et al., 2025), our results highlight genomic GC content as a crucial covarying factor. The similar evolutionary dynamics we observe between genome size and GC content parallel findings in microbes (Gao & Wu, 2022), suggesting this relationship represents a fundamental taxon-independent principle of genomic architecture evolution.

The strong phylogenetic signal of chromosome number across most orders suggests deep evolutionary conservation. Its evolution under a pulsed (PE1_BM) model indicates that this conserved diversity was generated during specific periods of rapid karyotypic change, after which chromosome numbers stabilized within lineages. The absence of phylogenetic signal in coleopteran chromosome number indicates either exceptionally high evolutionary lability or a sampling artifact. Furthermore, chromosome number showed no significant evolutionary relationship with either genome size or GC content in any order, indicating these major genomic traits evolve independently of karyotypic reorganization in insects. *Diptera* exhibits strong phylogenetic signal with a punctuated kappa model (κ = 0.595), while *Hymenoptera* shows minimal phylogenetic dependence—non-coding gene counts follow white noise evolution, and coding gene counts display no significant signal. Corresponding family-level analyses support these order-level findings, particularly confirming the absence of phylogenetic signals in *Hymenoptera*. For the remaining orders, limited sampling constrains robust inference into their evolutionary modes.

Critically, both morphological and genomic traits show no simple correlation with species richness when analyzed through traditional metrics like phylogenetic signal or by merely identifying the predominant evolutionary model. Genomic traits consistently point towards punctuated models, while morphological traits show no uniform association with either Brownian or pulsed modes. The key insight emerges not from whether pulses occur, but from their macroevolutionary pattern: species-poor lineages appear to exhibit pronounced pulses in morphological traits, whereas species-rich lineages channel major evolutionary jumps into genomic architecture. While more extensive taxonomic and phenotypic sampling is required to substantiate the morphological pattern—and to test whether these pulses represent maladaptive trajectories that contribute to clade reduction—our results highlight the need for mechanistic models that can explain how large-scale genomic pulses might subsequently influence lineage diversification. While phylogenetic signal shows no clear trend with species richness, the prevalence of kappa models for genome size and GC content in species-rich orders suggests a greater tendency for evolutionary change concentrated at speciation events (Gould & Eldredge, 1977), a dynamic which may itself contribute to their heightened diversification potential. We note that the strength of phylogenetic constraint for both morphological and genomic traits is independent of evolutionary distance between orders, revealing that trait evolutionary modes are labile properties that can arise in any lineage. Consequently, the phylogenetic signal of a trait is not a simple function of a lineage’s diversification history or its position in the tree of life.

These results raise the possibility that the macroevolutionary pattern of pulsed evolution holds greater significance than its mere presence. A tentative model emerges: morphological pulses in species-poor clades could reflect maladaptive trajectories, whereas species-rich clades may channel substantial evolutionary change into genomic traits, as seen in the genome size pulses of Diptera and Hymenoptera. Elucidating the mechanistic basis for this pattern represents a key objective for future research. The contrasting case of Lepidoptera, however, indicates this is not a universal rule.

In conclusion, our study reveals a fundamental decoupling in insect evolution: genomic architecture evolves through pulsed, diversification-independent processes, while morphological traits are shaped by developmental constraints that favor a more gradual mode of evolution. Critically, within this morphological landscape, we find that both evolutionary lability in certain traits and pulsed shifts in others can serve as potential engines for diversification. Quantifying these distinct macroevolutionary pathways provides the critical evolutionary foundation needed for accurate ecological modeling, ultimately enhancing our ability to predict functional diversity and species’ responses to environmental change —and offering new perspectives on the evolutionary underpinnings of insects’ ecological success.

## Data Availability Statement

All data and R scripts used in this study, including morphological and genomic datasets and analysis code for phylogenetic and random forest models, will be made publicly available upon acceptance of the manuscript. The data and scripts will be deposited in a public GitHub repository to ensure accessibility for further research and replication.

## Acknowledgements

The research infrastructure used in this study was supported by a grant from the Department of Biotechnology (DBT), Government of India [Grant Number: BT/PR59599/BID/7/1087/2025]. AA acknowledges the valuable discourse at INCOB2025 concerning evolutionary pattern convergence, which motivated enhanced comparative frameworks in this study.

## Declaration of AI-Assisted Writing Support

The authors acknowledge the use of generative AI during the preparation of this manuscript exclusively for language polishing, grammatical correction, and sentence structure refinement. All scientific content, data analysis, interpretation of results, and conceptual development were solely conducted by the human authors. The AI tool functioned as a writing assistant and did not contribute to any scientific reasoning, experimental design, data interpretation, or conceptual insights. The final manuscript has been thoroughly reviewed and approved by all authors, who take full responsibility for its scientific content and integrity

## References

Ackerly, David. “Conservatism and diversification of plant functional traits: evolutionary rates versus phylogenetic signal.” Proceedings of the National Academy of Sciences 106, no. supplement_2 (2009): 19699–19706.

Adamski, Zbigniew, Sabino A. Bufo, Szymon Chowański, Patrizia Falabella, Jan Lubawy, Paweł Marciniak, Joanna Pacholska-Bogalska et al. “Beetles as model organisms in physiological, biomedical and environmental studies–a review.” Frontiers in physiology 10 (2019): 319.

Aguiar, Alexandre P., Andrew R. Deans, Michael S. Engel, Mattias Forshage, John T. Huber, John T. Jennings, Norman F. Johnson et al. “Order Hymenoptera. In: Zhang, Z.-Q.(Ed.) Animal biodiversity: an outline of higher-level classification and survey of taxonomic richness (Addenda 2013).” Zootaxa 3703, no. 1 (2013): 51–62.

Ajay, Akash, Tina Begum, Ajay Arya, Krishan Kumar, and Shandar Ahmad. “Global and local genomic features together modulate the spontaneous single nucleotide mutation rate.” Computational Biology and Chemistry 112 (2024): 108107.

Akaike, Hirotugu. “Akaike’s information criterion.” In International encyclopedia of statistical science, pp. 41–42. Springer, Berlin, Heidelberg, 2025.

Alfsnes, Kristian, Hans Petter Leinaas, and Dag Olav Hessen. “Genome size in arthropods; different roles of phylogeny, habitat and life history in insects and crustaceans.” Ecology and evolution 7, no. 15 (2017): 5939–5947.

Arakawa, Kazuharu, and Masaru Tomita. “The GC skew index: a measure of genomic compositional asymmetry and the degree of replicational selection.” Evolutionary Bioinformatics 3 (2007): 117693430700300006.

Auman, Tzach, and Ariel D. Chipman. “The evolution of gene regulatory networks that define arthropod body plans.” Integrative and Comparative Biology 57, no. 3 (2017): 523–532.

Bardua, Carla, Mark Wilkinson, David J. Gower, Emma Sherratt, and Anjali Goswami. “Morphological evolution and modularity of the caecilian skull.” BMC evolutionary biology 19, no. 1 (2019): 30.

Beccaloni, George, and Paul Eggleton. “Order Blattodea. In: Zhang, Z.-Q.(Ed.) Animal Biodiversity: An Outline of Higher-level Classification and Survey of Taxonomic Richness (Addenda 2013).” Zootaxa 3703, no. 1 (2013): 46–48.

Berger, Vance W., and YanYan Zhou. “Kolmogorov–smirnov test: Overview.” Wiley statsref: Statistics reference online (2014).

Blomberg, Simon P., Theodore Garland Jr, and Anthony R. Ives. “Testing for phylogenetic signal in comparative data: behavioral traits are more labile.” Evolution 57, no. 4 (2003): 717–745.

Bruce, Toby JA. “Interplay between insects and plants: dynamic and complex interactions that have coevolved over millions of years but act in milliseconds.” Journal of Experimental Botany 66, no. 2 (2015): 455–465.

Buezas, Guido N., Federico Becerra, Alejandra I. Echeverría, Adrián Cisilino, and Aldo I. Vassallo. “Mandible strength and geometry in relation to bite force: a study in three caviomorph rodents.” Journal of Anatomy 234, no. 4 (2019): 564–575.

Bui, Van Bac, Thomas Ziegler, and Michael Bonkowski. “Morphological traits reflect dung beetle response to land use changes in tropical karst ecosystems of Vietnam.” Ecological Indicators 108 (2020): 105697.

Burnham, Kenneth P., and David R. Anderson, eds. Model selection and multimodel inference: a practical information-theoretic approach. New York, NY: Springer New York, 2002.

Cerritos, René. “Insects as food: an ecological, social and economical approach.” CABI Reviews 2009 (2009): 1-10.

Choi, Ik-Young, Eun-Chae Kwon, and Nam-Soo Kim. “The C-and G-value paradox with polyploidy, repeatomes, introns, phenomes and cell economy.” Genes & genomics 42, no. 7 (2020): 699–714.

Christiansen, P. “Evolutionary implications of bite mechanics and feeding ecology in bears.” Journal of Zoology 272, no. 4 (2007): 423–443.

Christiansen, Per, and Stephen Wroe. “Bite forces and evolutionary adaptations to feeding ecology in carnivores.” Ecology 88, no. 2 (2007): 347–358.

Cigliano, María Marta, Holger Braun, David C. Eades, and Daniel Otte. “Orthoptera species file.” (2024).

Cunha, Marina Souza, Danon Clemes Cardoso, Maykon Passos Cristiano, Lucio Antônio de Oliveira Campos, and Denilce Meneses Lopes. “The bee chromosome database (Hymenoptera: Apidae).” Apidologie 52, no. 2 (2021): 493–502.

dos Santos-Pinto, José Roberto Aparecido, Amilcar Perez-Riverol, Alexis Musacchio Lasa, and Mario Sergio Palma. “Diversity of peptidic and proteinaceous toxins from social Hymenoptera venoms.” Toxicon 148 (2018): 172–196.

Ding, Changping, and Yalin Zhang. “Phylogenetic relationships of Pieridae (Lepidoptera: Papilionoidea) in China based on seven gene fragments.” Entomological Science 20, no. 1 (2017): 15–23.

Dudley, Robert. The biomechanics of insect flight: form, function, evolution. Princeton university press, 2002.

Duncan, Mataya, Michael DeGiorgio, Raquel Assis, and Richard Adams. “Robust regression rescues poor phylogenetic decisions.” BMC Ecology and Evolution 25, no. 1 (2025): 108.

Edel, Carina, Peter T. Rühr, Melina Frenzel, Thomas van de Kamp, Tomáš Faragó, Jörg U. Hammel, Fabian Wilde, and Alexander Blanke. “Bite force transmission and mandible shape in grasshoppers, crickets, and allies is not driven by dietary niches.” Evolution 78, no. 12 (2024): 1958–1968.

Engel, Michael S. “Insect evolution.” Current Biology 25, no. 19 (2015): R868–R872.

Espeland, Marianne, Jason PW Hall, Philip J. DeVries, David C. Lees, Mark Cornwall, Yu-Feng Hsu, Li-Wei Wu et al. “Ancient Neotropical origin and recent recolonisation: Phylogeny, biogeography and diversification of the Riodinidae (Lepidoptera: Papilionoidea).” Molecular phylogenetics and evolution 93 (2015): 296–306.

Felice, Ryan N., Akinobu Watanabe, Andrew R. Cuff, Eve Noirault, Diego Pol, Lawrence M. Witmer, Mark A. Norell, Patrick M. O’Connor, and Anjali Goswami. “Evolutionary integration and modularity in the archosaur cranium.” Integrative and comparative biology 59, no. 2 (2019): 371–382.

Felsenstein, J., 1985. Phylogenies and the comparative method. The American Naturalist, 125(1), pp.1–15.

Foottit, R. G., HEL V. Maw, C. D. Von Dohlen, and P. D. N. Hebert. “Species identification of aphids (Insecta: Hemiptera: Aphididae) through DNA barcodes.” Molecular Ecology Resources 8, no. 6 (2008): 1189–1201.

Foster, Woodbridge A., and Edward D. Walker. “Chapter 15-Mosquitoes (Culicidae).” Medical and veterinary entomology 2019 (2019): 261–325.

FitzJohn, Richard G. “Diversitree: comparative phylogenetic analyses of diversification in R.” Methods in ecology and evolution 3, no. 6 (2012): 1084–1092.

Gao, Y., & Wu, M. (2022). Microbial genomic trait evolution is dominated by frequent and rare pulsed evolution. Science Advances, 8(28), eabn1916.

Garland Jr, Theodore, Paul H. Harvey, and Anthony R. Ives. “Procedures for the analysis of comparative data using phylogenetically independent contrasts.” Systematic biology 41, no. 1 (1992): 18–32.

Gould, Stephen Jay, and Niles Eldredge. “Punctuated equilibria: the tempo and mode of evolution reconsidered.” Paleobiology 3, no. 2 (1977): 115–151.

Graça, Márlon Breno, and M. Alma Solis. “Order Lepidoptera.” In Thorp and Covich’s Freshwater Invertebrates, pp. 325–338. Academic Press, 2018.

Graphodatsky, Alexander S., Vladimir A. Trifonov, and Roscoe Stanyon. “The genome diversity and karyotype evolution of mammals.” Molecular cytogenetics 4, no. 1 (2011): 22.

Gross, Juergen, Uwe Ligges, and Maintainer Uwe Ligges. “Package ‘nortest’.” Five omnibus tests for testing the composite hypothesis of normality (2015).

Guerra, Mꎬ. “Chromosome numbers in plant cytotaxonomy: concepts and implications.” Cytogenetic and Genome research 120, no. 3-4 (2008): 339–350.

Hagge, Jonas, Jörg Müller, Tone Birkemoe, Jörn Buse, Martin M. Gossner, Axel Gruppe, Christoph Heibl et al. “What does a threatened saproxylic beetle look like? Modelling extinction risk using a new morphological trait database.” Journal of Animal Ecology 90, no. 8 (2021): 1934–1947.

Hansen, Thomas F. “Stabilizing selection and the comparative analysis of adaptation.” Evolution 51, no. 5 (1997): 1341–1351.

Harmon, Luke J., Jonathan B. Losos, T. Jonathan Davies, Rosemary G. Gillespie, John L. Gittleman, W. Bryan Jennings, Kenneth H. Kozak et al. “Early bursts of body size and shape evolution are rare in comparative data.” Evolution 64, no. 8 (2010): 2385–2396.

Harmon, Luke, Jason Weir, Chad Brock, Rich Glor, Wendell Challenger, Gene Hunt, Rich FitzJohn et al. “Package ‘geiger’.” R package version 2, no. 3 (2015): 1–74.

Heckenhauer, Jacqueline, Paul B. Frandsen, John S. Sproul, Zheng Li, Juraj Paule, Amanda M. Larracuente, Peter J. Maughan, et al. “Genome size evolution in the diverse insect order Trichoptera.” GigaScience 11 (2022): giac011.

Heppner, John B. “Geometer Moths (Lepidoptera: Geometridae).” In Encyclopedia of Entomology, pp. 1610–1611. Springer, Dordrecht, 2008.

Herbert, Mélanie CM, André Nel, Brian V. Brown, Antonio Arillo, Brendon E. Boudinot, and Mónica M. Solórzano-Kraemer. “Review of wing morphology in fossil and modern species of humpbacked flies (Diptera: Phoridae).” BMC biology 23, no. 1 (2025): 298.

Huang, Weidong, Changhua Zhang, Tingzhen Zhang, Ye Xu, Shiwen Xu, Li Tian, Hu Li, Wanzhi Cai, and Fan Song. “Features and evolution of control regions in leafroller moths (Lepidoptera: Tortricidae) inferred from mitochondrial genomes and phylogeny.” International Journal of Biological Macromolecules 236 (2023): 123928.

Huber, John T. “Biodiversity of hymenoptera.” Insect biodiversity: science and society (2017): 419–461.

Hunt, Gene. “Gradual or pulsed evolution: when should punctuational explanations be preferred?.” Paleobiology 34, no. 3 (2008): 360–377.

Isip, Justin E., Marc EH Jones, and Natalie Cooper. “Clade-wide variation in bite-force performance is determined primarily by size, not ecology.” Proceedings of the Royal Society B 289, no. 1969 (2022): 20212493.

Joop, Gerrit, and Andreas Vilcinskas. “Coevolution of parasitic fungi and insect hosts.” Zoology 119, no. 4 (2016): 350–358.

Jolivet, Pierre, Eduard Petitpierre, and Ting H. Hsiao, eds. Biology of chrysomelidae. Vol. 42. Springer Science & Business Media, 2012.

Kalkman, Vincent J., Viola Clausnitzer, Klaas-Douwe B. Dijkstra, Albert G. Orr, Dennis R. Paulson, and Jan Van Tol. “Global diversity of dragonflies (Odonata) in freshwater.” Hydrobiologia 595, no. 1 (2008): 351–363.

Keegan, Kevin L., Jadranka Rota, Reza Zahiri, Alberto Zilli, Niklas Wahlberg, B. Christian Schmidt, J. Donald Lafontaine, Paul Z. Goldstein, and David L. Wagner. “Toward a stable global Noctuidae (Lepidoptera) taxonomy.” Insect Systematics and Diversity 5, no. 3 (2021): 1.

Kristensen, Niels P., Malcolm J. Scoble, and O. L. E. Karsholt. “Lepidoptera phylogeny and systematics: the state of inventorying moth and butterfly diversity.” Zootaxa 1668, no. 1 (2007): 699–747.

Kyriacou, Riccardo G., Peter O. Mulhair, and Peter WH Holland. “GC content across insect genomes: phylogenetic patterns, causes and consequences.” Journal of Molecular Evolution 92, no. 2 (2024): 138–152.

Li, Daijiang, Russell Dinnage, Lucas A. Nell, Matthew R. Helmus, and Anthony R. Ives. “phyr: An r package for phylogenetic species-distribution modelling in ecological communities.” Methods in Ecology and Evolution 11, no. 11 (2020): 1455–1463.

Li, Fei, Xianxin Zhao, Meizhen Li, Kang He, Cong Huang, Yuenan Zhou, Zekai Li, and James R. Walters. “Insect genomes: progress and challenges.” Insect molecular biology 28, no. 6 (2019): 739–758.

Li, Xinghao, Rufan Li, Fuqiang Rao, Rong An, Jianchang Li, Zhenlan Zhang, Yonghong Li, and Deguang Liu. “Genetic characterization of 2 Ceutorhynchus (Coleoptera: Curculionidae) weevils with mitogenomes and insights into the phylogeny and evolution of related weevils.” Journal of Insect Science 24, no. 2 (2024): 14.

Losey, John E., and Mace Vaughan. “The economic value of ecological services provided by insects.” Bioscience 56, no. 4 (2006): 311–323.

Losos, Jonathan B. “Phylogenetic niche conservatism, phylogenetic signal and the relationship between phylogenetic relatedness and ecological similarity among species.” Ecology letters 11, no. 10 (2008): 995–1003.

Madi, Naïma, Michiel Vos, Carmen Lia Murall, Pierre Legendre, and B. Jesse Shapiro. “Does diversity beget diversity in microbiomes?.” Elife 9 (2020): e58999.

Mattick, John S. “Non-coding RNAs: the architects of eukaryotic complexity.” EMBO reports (2001).

Mayhew, Peter J. “Comparative analysis of behavioural traits in insects.” Current opinion in insect science 27 (2018): 52–60.

McKenna, Duane D., and Brian D. Farrell. “Beetles (Coleoptera).” The timetree of life 278 (2009): 289.

Méneville, G. A., J. B. Robineau-Desvoidy, P. J. M. Macquart, C. H. Blanchard, J. A. Laboulbène, J. M. F. Bigot, J. O. Westwood et al. “Neotropical Diptera.”

Misof, Bernhard, Shanlin Liu, Karen Meusemann, Ralph S. Peters, Alexander Donath, Christoph Mayer, Paul B. Frandsen et al. “Phylogenomics resolves the timing and pattern of insect evolution.” Science 346, no. 6210 (2014): 763–767.

Molina-Venegas, Rafael, and Miguel Á. Rodríguez. “Revisiting phylogenetic signal; strong or negligible impacts of polytomies and branch length information?.” BMC evolutionary biology 17, no. 1 (2017): 53.

Münkemüller, Tamara, Sébastien Lavergne, Bruno Bzeznik, Stéphane Dray, Thibaut Jombart, Katja Schiffers, and Wilfried Thuiller. “How to measure and test phylogenetic signal.” Methods in Ecology and Evolution 3, no. 4 (2012): 743–756.

Nijhout, H. Frederik, and Viviane Callier. “Developmental mechanisms of body size and wing-body scaling in insects.” Annual Review of Entomology 60, no. 1 (2015): 141–156.

Nijhout, H. Frederik, Lynn M. Riddiford, Christen Mirth, Alexander W. Shingleton, Yuichiro Suzuki, and Viviane Callier. “The developmental control of size in insects.” Wiley Interdisciplinary Reviews: Developmental Biology 3, no. 1 (2014): 113–134.

Oliveira, Marisa M., Alexander W. Shingleton, and Christen K. Mirth. “Coordination of wing and whole-body development at developmental milestones ensures robustness against environmental and physiological perturbations.” PLoS genetics 10, no. 6 (2014): e1004408.

Ospina-Garcés, Sandra M., Federico Escobar, Martha L. Baena, Adrian LV Davis, and Clarke H. Scholtz. “Do dung beetles show interrelated evolutionary trends in wing morphology, flight biomechanics and habitat preference?.” Evolutionary Ecology 32, no. 6 (2018): 663–682.

Orme, David, Rob Freckleton, Gavin Thomas, Thomas Petzoldt, Susanne Fritz, Nick Isaac, and Will Pearse. “The caper package: comparative analysis of phylogenetics and evolution in R.” R package version 5, no. 2 (2013): 1–36.

Pagel, Mark, Chris Venditti, and Andrew Meade. “Large punctuational contribution of speciation to evolutionary divergence at the molecular level.” Science 314, no. 5796 (2006): 119–121.

Pagel, Mark. “The maximum likelihood approach to reconstructing ancestral character states of discrete characters on phylogenies.” Systematic biology 48, no. 3 (1999): 612–622.

Paradis, Emmanuel, Simon Blomberg, Ben Bolker, Joseph Brown, Julien Claude, Hoa Sien Cuong, Richard Desper, and Gilles Didier. “Package ‘ape’.” Analyses of phylogenetics and evolution, version 2, no. 4 (2019): 47.

Pennell, Matthew W., Jonathan M. Eastman, Graham J. Slater, Joseph W. Brown, Josef C. Uyeda, Richard G. FitzJohn, Michael E. Alfaro, and Luke J. Harmon. “geiger v2. 0: an expanded suite of methods for fitting macroevolutionary models to phylogenetic trees.” Bioinformatics 30, no. 15 (2014): 2216–2218.

Petrov, Dmitri A. “Evolution of genome size: new approaches to an old problem.” TRENDS in Genetics 17, no. 1 (2001): 23–28.

Porto, Arthur, Felipe B. de Oliveira, Leila T. Shirai, Valderes De Conto, and Gabriel Marroig. “The evolution of modularity in the mammalian skull I: morphological integration patterns and magnitudes.” Evolutionary Biology 36, no. 1 (2009): 118–135.

Resh, Vincent H., and Ring T. Cardé, eds. Encyclopedia of insects. Academic press, 2009.

Revell, Liam J. “phytools 2.0: an updated R ecosystem for phylogenetic comparative methods (and other things).” PeerJ 12 (2024): e16505.

Ricklefs, Robert E. “Reconciling diversification: Random pulse models of speciation and extinction.” The American Naturalist 184, no. 2 (2014): 268–276.

Ryan Gregory, T. “Genome size and developmental complexity.” Genetica 115, no. 1 (2002): 131–146.

Saif, Nasif Bin, Ramin JA Guilani, Shayan Ramezanpour, Arman Toofani, Sepehr H. Eraghi, Geoff Goss, Chung-Ping Lin, Stanislav Gorb, and Hamed Rajabi. “Engineering the battle: Design-specific analysis of stag beetle mandibles for combat efficiency.” PNAS Nexus 4, no. 7 (2025): pgaf205.

Salman, Muhammad, Ping Wang, Nian Liu, Xuanzeng Liu, Weian Deng, and Yuan Huang. “Evolutionary dynamics of repetitive elements and genome size in Tetrigidae (Orthoptera: Caelifera).” Scientific Reports 15, no. 1 (2025): 34509.

Santana, Sharlene E., Elizabeth R. Dumont, and Julian L. Davis. “Mechanics of bite force production and its relationship to diet in bats.” Functional Ecology 24, no. 4 (2010): 776–784.

Scholz, Fritz W., and Michael A. Stephens. “K-sample Anderson–Darling tests.” Journal of the American Statistical Association 82, no. 399 (1987): 918–924.

Simon, Sabrina, Harald Letsch, Sarah Bank, Thomas R. Buckley, Alexander Donath, Shanlin Liu, Ryuichiro Machida et al. “Old World and New World Phasmatodea: phylogenomics resolve the evolutionary history of stick and leaf insects.” Frontiers in Ecology and Evolution 7 (2019): 345.

Shatkovska, O. V., and M. Ghazali. “Relationship between developmental modes, flight styles, and wing morphology in birds.” The European Zoological Journal 84, no. 1 (2017): 390–401.

Song, Hojun, and Sibyl R. Bucheli. “Comparison of phylogenetic signal between male genitalia and non-genital characters in insect systematics.” Cladistics 26, no. 1 (2010): 23–35.

Symonds, Matthew RE, and Simon P. Blomberg. “A primer on phylogenetic generalised least squares.” In Modern phylogenetic comparative methods and their application in evolutionary biology: concepts and practice, pp. 105–130. Berlin, Heidelberg: Springer Berlin Heidelberg, 2014.

Tan, Wei Cheng, John Measey, Bieke Vanhooydonck, and Anthony Herrel. “The relationship between bite force, morphology, and diet in southern African agamids.” BMC Ecology and Evolution 21, no. 1 (2021): 126.

Thomas, Gregg WC, Elias Dohmen, Daniel ST Hughes, Shwetha C. Murali, Monica Poelchau, Karl Glastad, Clare A. Anstead et al. “Gene content evolution in the arthropods.” Genome biology 21, no. 1 (2020): 15.

Waage, Jeffrey K. “The evolution of insect/vertebrate associations.” Biological journal of the Linnean Society 12, no. 3 (1979): 187–224.

Wang, Yiguan, and Darren J. Obbard. “Experimental estimates of germline mutation rate in eukaryotes: a phylogenetic meta-analysis.” Evolution Letters 7, no. 4 (2023): 216–226.

Whittaker, Robert J., Richard J. Ladle, Miguel B. Araújo, José María Fernández-Palacios, Juan Domingo Delgado, and José Ramón Arévalo. “The island immaturity–speciation pulse model of island evolution: an alternative to the “diversity begets diversity” model.” Ecography 30, no. 3 (2007): 321–327.

Wiens, John J. “The niche, biogeography and species interactions.” Philosophical Transactions of the Royal Society B: Biological Sciences 366, no. 1576 (2011): 2336–2350.

Wilson, Edward O., and James Turner. “Order: Hemiptera.” Arthropod Fauna of the United Arab Emirates 2010 (2010): 113–125.

Yan, Zhen-Tian, Zhen-Huai Fan, Shu-Lin He, Xue-Qian Wang, Bin Chen, and Si-Te Luo. “Mitogenomes of eight Nymphalidae butterfly species and reconstructed phylogeny of Nymphalidae (Nymphalidae: Lepidoptera).” Genes 14, no. 5 (2023): 1018.

Yan, Zhen-Tian, Xiao-Ya Tang, Dong Yang, Zhen-Huai Fan, Si-Te Luo, and Bin Chen. “Phylogenetic and comparative genomics study of Papilionidae based on mitochondrial genomes.” Genes 15, no. 7 (2024): 964.

Yuan, Hao, Xiao-Jing Liu, Xuan-Zeng Liu, Li-Na Zhao, Shao-Li Mao, and Yuan Huang. “The evolutionary dynamics of genome sizes and repetitive elements in Ensifera (Insecta: Orthoptera).” BMC genomics 25, no. 1 (2024): 1041.

Yuan, Hao, Yuan Huang, Ying Mao, Nan Zhang, Yimeng Nie, Xue Zhang, Yafu Zhou, and Shaoli Mao. “The evolutionary patterns of genome size in Ensifera (Insecta: Orthoptera).” Frontiers in Genetics 12 (2021): 693541.

Zhang, Hong-Li, and Fei Ye. “Comparative mitogenomic analyses of praying mantises (Dictyoptera, Mantodea): origin and evolution of unusual intergenic gaps.” International Journal of Biological Sciences 13, no. 3 (2017): 367.

