## Supplementary Figures for "The Duality of Insect Macroevolution: Pulsed Genomes and Gradual Morphology Shape Lineage Diversification"

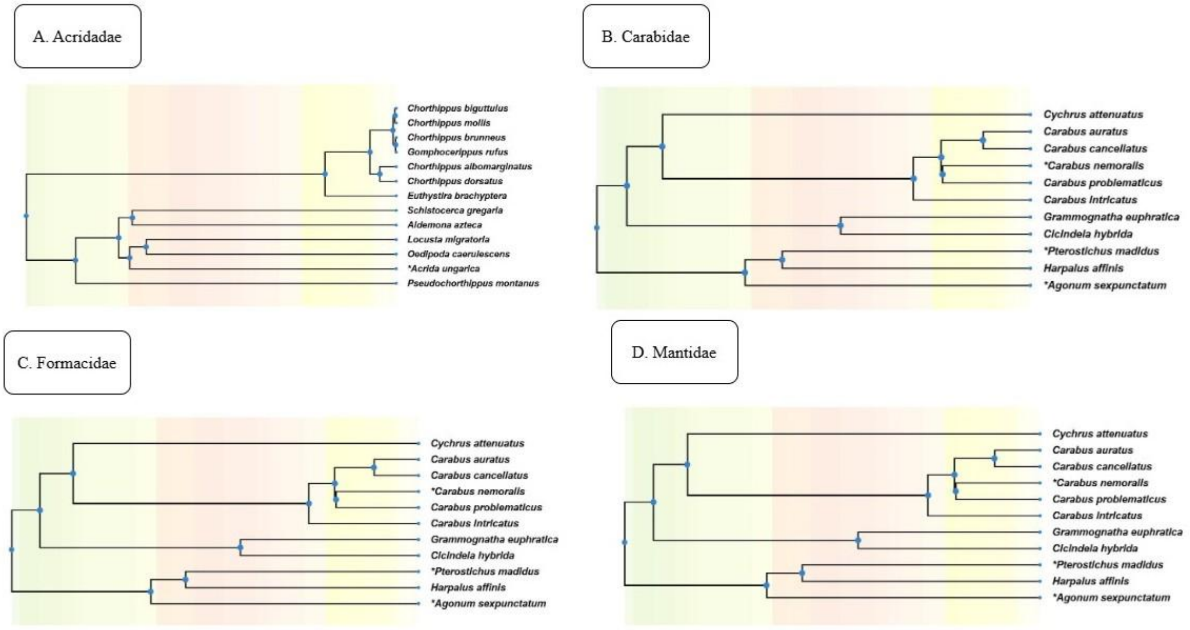


**Supplementary Figure 1 (SF1)**: Representative Family-Level Phylogenies for Morphological Trait Evolution.

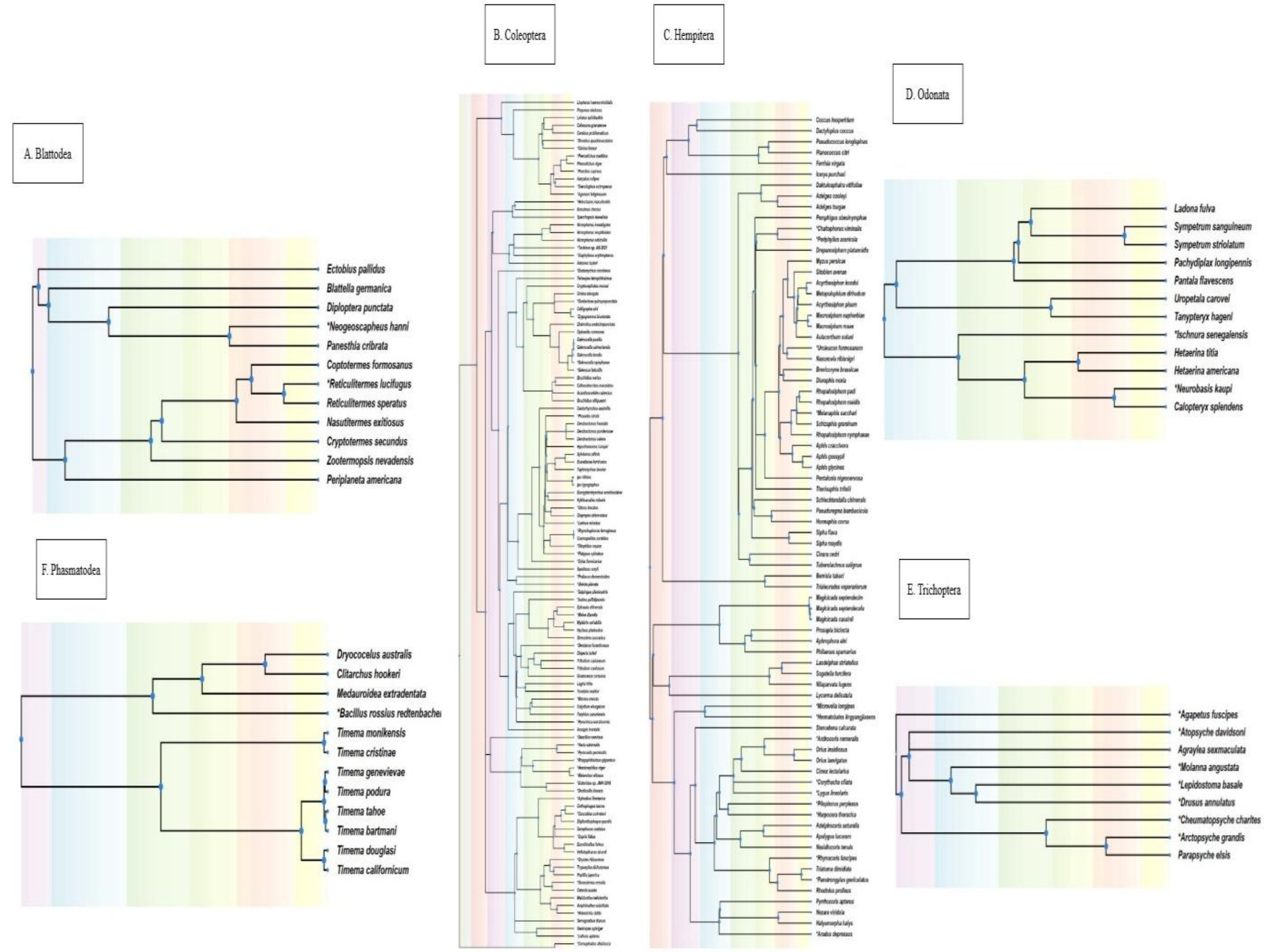


**Supplementary Figure 2 (SF2):** Representative Family-Level Phylogenies for Genomic Trait Evolution.

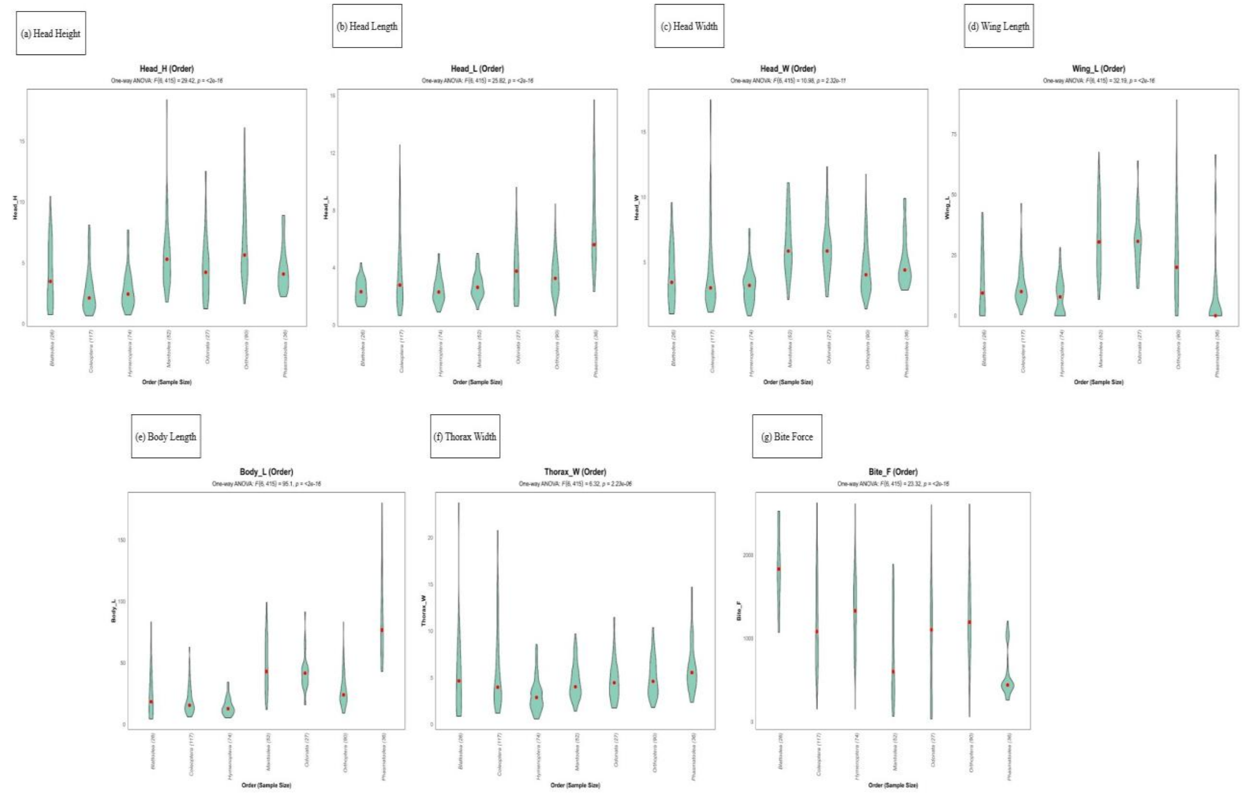


**Supplementary Figure 3 (SF3).** Distribution and ANOVA of Morphological Traits Across Insect Orders.

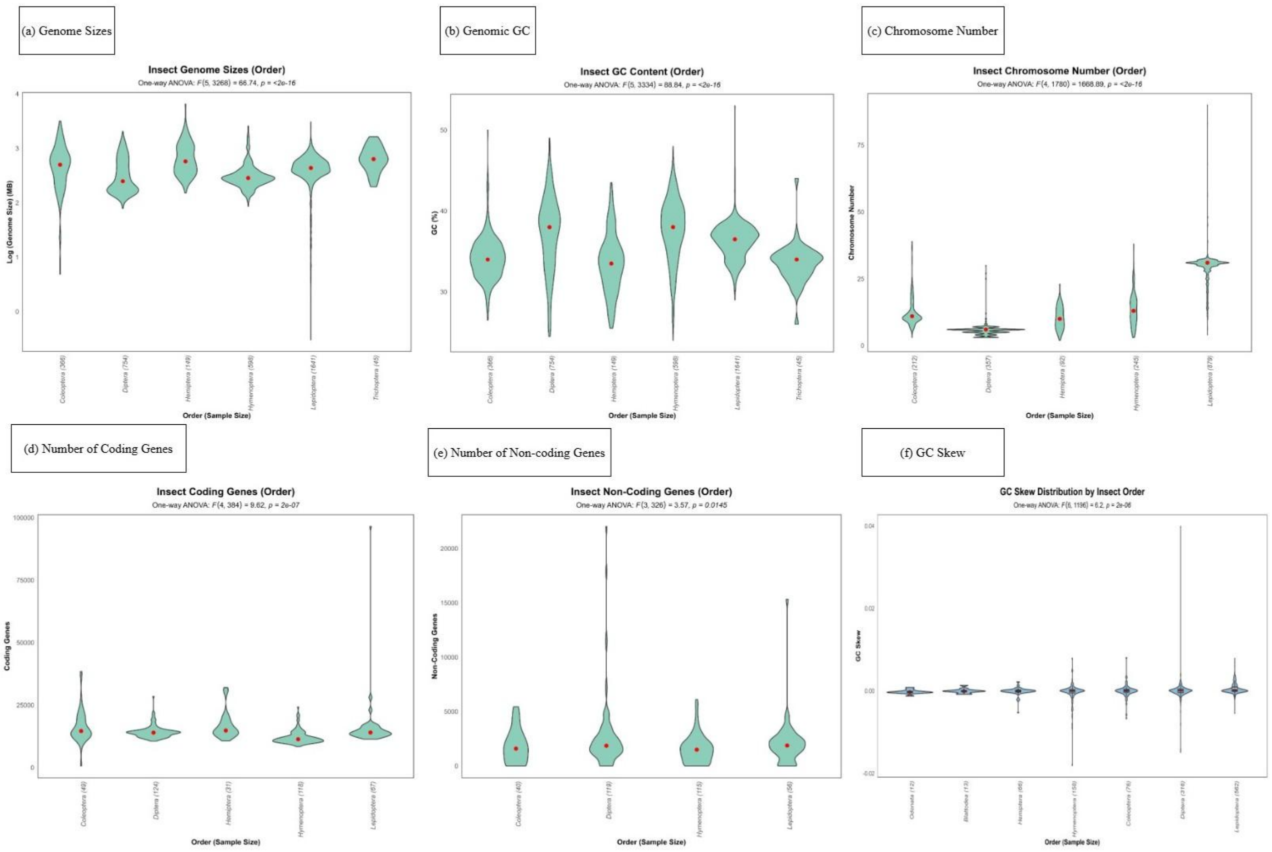


**Supplementary Figure 4 (SF4).** Distribution and ANOVA of Morphological Traits Across Insect Orders.

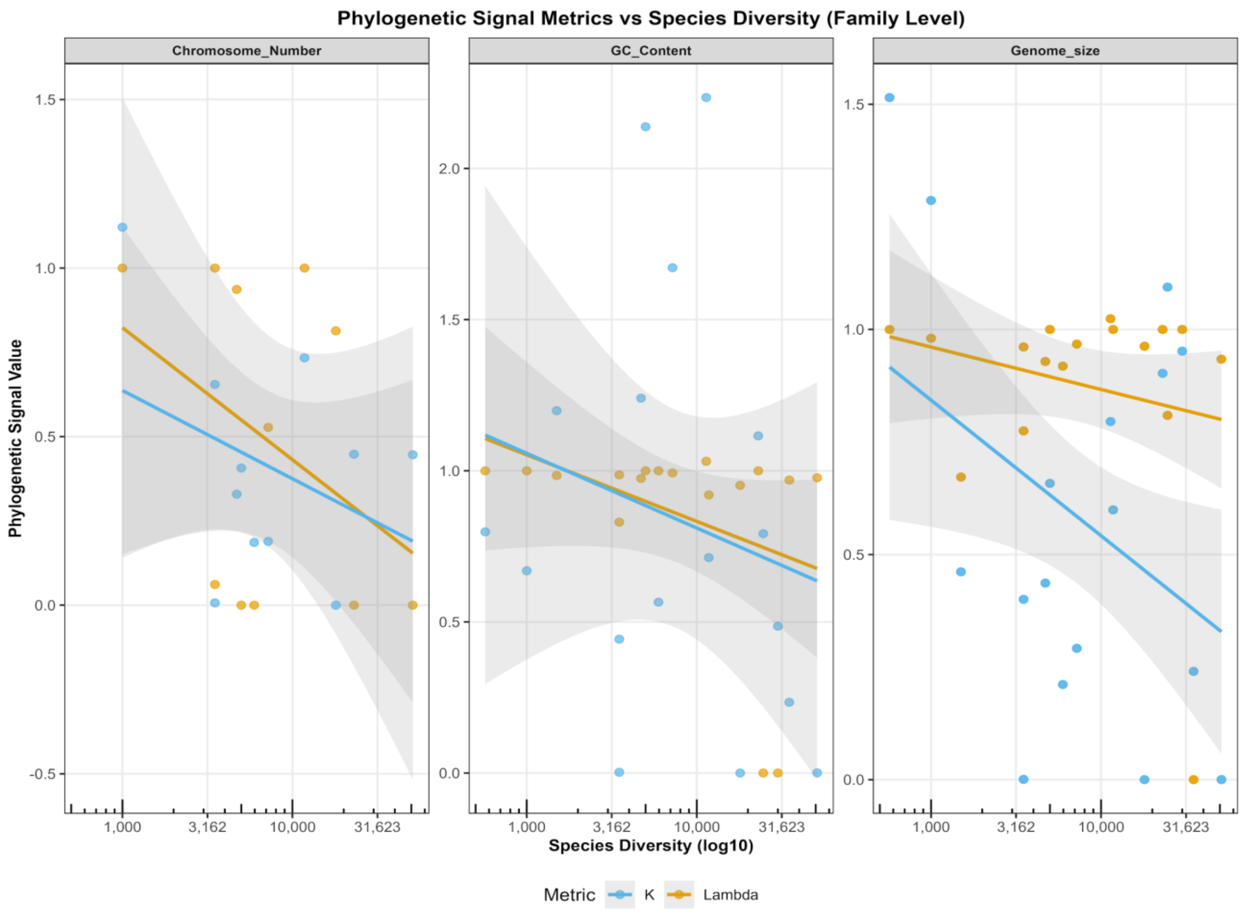


**Supplementary Figure 5 (SF5).** Relationship Between Species Richness and Phylogenetic Signal of Genomic Traits.
